# Beyond morphometry: Adolescent brain microarchitecture reflects sex differences in pubertal timing and hormones

**DOI:** 10.64898/2026.11.17.766192

**Authors:** Erynn Christensen, Elvisha Dhamala, Megan M. Herting, Katherine L. Bottenhorn

## Abstract

Puberty is a sensitive neurodevelopmental period characterized by dramatic increases in adrenal and gonadal hormones, yet how these changes relate to human brain microarchitecture remains unclear. Here, we used restriction spectrum imaging (RSI) to link peripubertal hormone levels to individual differences in brain microarchitecture (i.e., cellular and neurite density) in ∼5,000 peripubertal youth from the ABCD Study. Sex-and age-stratified multi-task regression analyses identified microarchitecture patterns jointly associated with physical pubertal maturation and each dehydroepiandrosterone (DHEA), testosterone, estradiol (in female youth), in turn, combining repeated cross-validation with permutation-based feature importance. At ages 9 to 11 years, microarchitecture differences explained 11-13% of pubertal hormone variability in female youth and 4-6% in male youth. Further, microarchitecture features were differentially associated with pubertal hormones. Younger male youth exhibited widespread pubertal hormone-related differences in neurite density that consolidated into fewer, stronger associations with age, while female youth exhibited age-related expansion in pubertal cellular density differences. Anatomically, hormone-relevant microarchitecture consistently included reward processing hubs (e.g., pallidum, accumbens, orbitofrontal cortex). Because animal models suggest that DHEA and testosterone metabolites have opposing effects on GABAergic signaling, we hypothesize that future research should investigate gray matter microarchitecture for insight into pubertal hormone contributions to adolescent refinement of excitatory-inhibitory balance. Ultimately, these findings provide a starting point for investigating how pubertal endocrine changes shape reward circuitry and neurotransmitter systems, with potential implications for the pubertal emergence of sex differences in adolescent psychopathology.

## Introduction

Adolescence spans a developmental period during which sex hormones exert permanent organizational effects on the brain (Larsen & Luna, 2018; Romeo, 2003; Schulz et al., 2009; Schulz & Sisk, 2016). Puberty is characterized by dramatic increases in sex hormones that cross the blood-brain-barrier and act on receptor-dense brain regions, including the prefrontal cortex, amygdala, and hippocampus (Frederiksen et al., 2024; Zsido et al., 2017). Animal studies demonstrate that testosterone and estradiol modulate cellular microarchitecture including dendritic spine density, synaptic pruning, neuron survival, and myelination (Juraska et al., 2013; McCarthy, 2008). In humans, neuroimaging studies have linked pubertal hormones to changes in gray and white matter morphometry, functional connectivity, and network development (Campbell et al., 2022a; Curtis et al., 2024; Herting et al., 2014; Khetan et al., 2026; Kovacs-Balint et al., 2023; Ladouceur et al., 2019). However, a significant gap remains in translating mechanistic, cellular insights to human neurodevelopment, which is critical for understanding how pubertal neuroendocrine processes contribute to the emergence of adolescent psychopathology.

Animal models link pubertal sex hormone increases to sexual differentiation of medial prefrontal cortex, amygdala, visual cortex, and hypothalamus (Juraska et al., 2013). Testosterone drives dendritic spine remodeling in rodent somatosensory cortex (Chen et al., 2013) and limbic system (Cunningham et al., 2007), regulates hippocampal neuron survival (Kight & McCarthy, 2020), and modulates amygdala cell death (De Lorme et al., 2012). These effects occur via androgen receptors, which are highly expressed in the human amygdala, primary visual cortex, striatum, and posterior superior temporal gyrus during early adolescence (Figure 1, first row). Organizational effects have also been linked to the absence of perinatal testosterone and the presence of ovarian hormones during puberty (Juraska et al., 2013). Estradiol regulates dendritic spine density, synaptogenesis, apoptosis, and neurite growth differently across brain regions (McCarthy, 2008); its primary central nervous system receptor, ER2β, is highly expressed in human medial prefrontal cortex, striatum, and posterior superior temporal gyrus (Figure 1, second row). Finally, dehydroepiandrosterone (DHEA) increases earlier in preadolescence with the activation of the hypothalamic-pituitary-adrenal (HPA) axis during adrenarche (Havelock et al., 2004), binding to both γ-aminobutyric acid (GABA_Α_) and N-methyl-D-aspartate (NMDA) receptors (Figure 1, third and fourth rows) (Compagnone & Mellon, 1998). Despite this wealth of mechanistic evidence from animal models, their implications for human microarchitecture development remain unclear.

**Figure 1.**
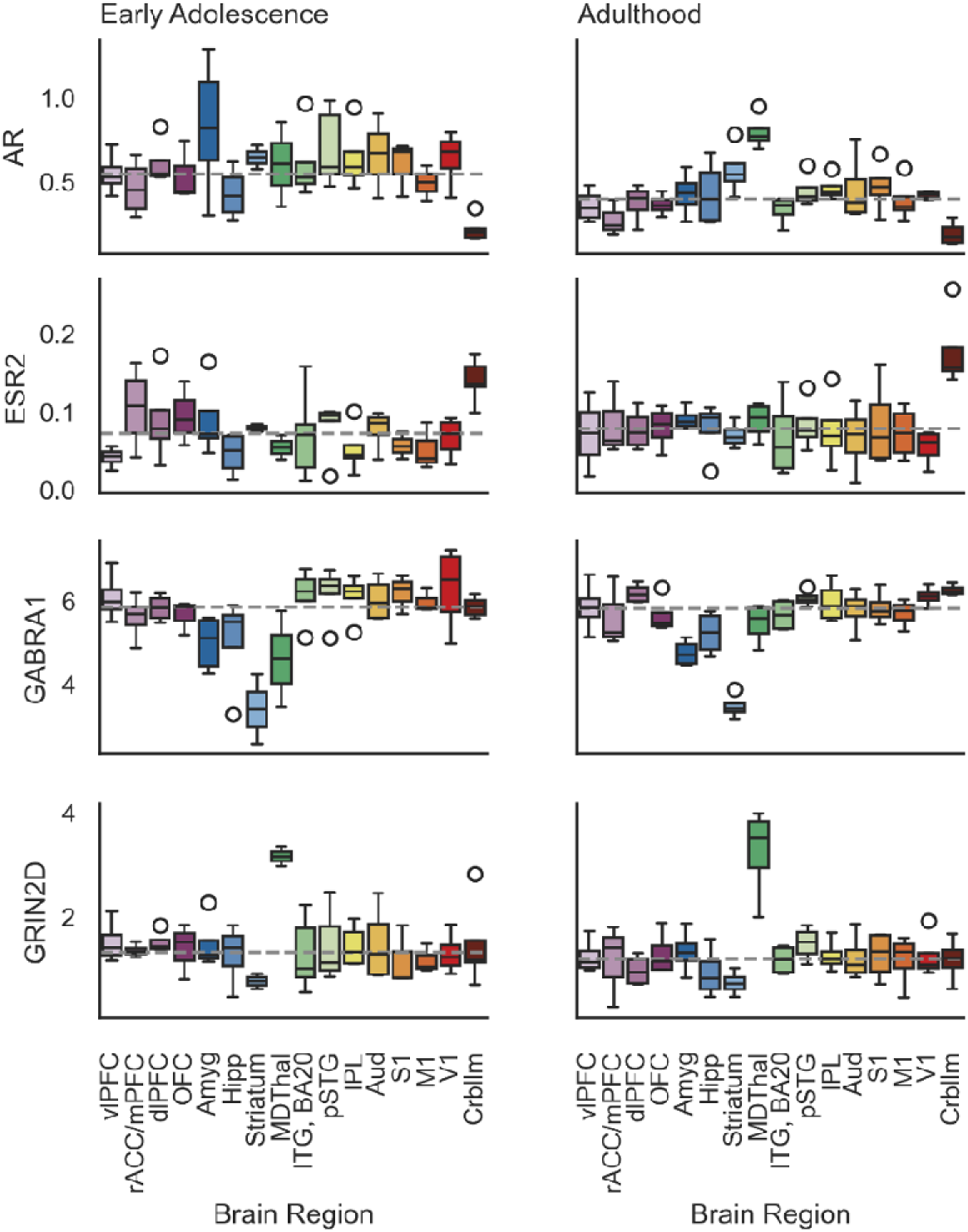
Hormone receptor expression across the brain. From the BrainSpan atlas of the developing human brain’s Developmental Transcriptome, expression of androgen receptors (AR, top row), estrogen receptor 2 (i.e., ERβ; ESR2, second row), γ-aminobutyric acid (GABA) A receptor alpha 1 (GABRA1, third row), glutamate receptor, ionotropic, N-methyl D-aspartate 2D (GRIN2D, fourth row) in early adolescents (ages 8 to 15 years, left) and adults (ages 24 to 60 years, right).

Diffusion-weighted magnetic resonance imaging (dMRI) may provide a means to bridge mechanistic research with the extant human neuroimaging literature, which focuses largely on gross morphometric measures from structural MRI scans. For example, cortical thickness and subcortical gray matter volumes measured from T1-weighted MRI scans can capture cell death and dendritic spine density (Rakic, 2009) and have been used to study hormone-related gray matter changes during puberty (Byrne et al., 2017; Curtis et al., 2024; Herting & Sowell, 2017; Khetan et al., 2026). In early adolescence, testosterone has been linked to changes in caudate and amygdala volume; estradiol, to overall gray and white matter volume (Campbell et al., 2022a; Herting et al., 2014). Restriction spectrum imaging (RSI) is a histologically validated dMRI approach that provides greater precision by separately modeling intra-and intercellular water diffusion (White et al., 2012, 2014). RSI further separates intracellular (i.e., restricted) diffusion into an isotropic component (RNI; reflecting neuronal or glial cell bodies) and a directional component (RND; reflecting axons and dendrites, collectively “neurites”, and glial processes). By estimating cellular and neurite density, RNI and RND may provide more precise insight into the role of hormones in pubertal brain development.

Human neuroimaging studies of adolescent development have been further limited by the measurement of challenges. Phasic (e.g., diurnal, seasonal) hormone fluctuations can obfuscate the sustained, tonic increases that drive lasting, organizational brain changes (Ankarberg & Norjavaara, 1999; Klusmann et al., 2024; Schulz et al., 2009). Conversely, physical markers of puberty (e.g., growth spurts, acne, body hair) measured via Tanner staging (Tanner, 1981) or the Pubertal Development Scale (PDS) (Petersen et al., 1988) are susceptible to subjective bias and lack detail during early puberty (Cheng et al., 2021). Because hormone levels and PDS scores present unique information about pubertal development, with complementary numerical properties, we can leverage their joint information to investigate how brain microarchitecture reflects individual differences in pubertal hormones. Regularized multi-task regression leverages this joint information to identify features that are related to both outcomes (Tabarestani et al., 2020), in this case brain regions that exhibit differences related to both the external, physical presentation of pubertal development from PDS scores and the internal, endocrine presentation of pubertal development from hormone levels. Here, we used regularized multi-task regression to leverage the shared variance between hormone levels and PDS scores and investigate individual differences in brain microarchitecture that reflect tonic, pubertal development.

We applied this modeling approach to data from the Adolescent Brain Cognitive Development Study (ABCD Study®) (Volkow et al., 2018), including 4,985 youth ages 9–11 years and 3,317 youth ages 11-13 years. Specifically, we identified brain microarchitecture related to pubertal DHEA and testosterone levels separately for female and male youth, and estradiol levels in female youth only, at 9-11 years of age (i.e., pre-early puberty) and 11-13 years of age (i.e., mid puberty). We expected pubertal DHEA-related microarchitecture differences in regions with high expression of GABA and NMDA receptors, including inferior and superior temporal gyri, ventrolateral prefrontal cortex, and medial thalamus. Likewise, we expected pubertal testosterone-related microarchitecture differences in regions with high ER2β expression (e.g., amygdala, primary visual cortex, inferior parietal lobule) and pubertal estrogen-related microarchitecture differences in regions with high ER2β expression (e.g., cerebellum, ventromedial prefrontal & orbitofrontal cortices). Across sexes and hormones, we expected to see stronger associations in mid puberty, when both hormone levels and neurodevelopmental trajectories exhibit greater variability compared to pre-/early puberty (Bottenhorn et al., 2023). Ultimately, identifying these sex-stratified, regional microarchitecture differences across early adolescence provides a preliminary neurobiological foundation for understanding how pubertal timing shapes adolescent brain maturation and vulnerability to psychopathology.

## Methods

Data used here come from the ongoing, longitudinal Adolescent Brain and Cognitive Development (ABCD) Study annual 5.1 data release (https://dx.doi.org/10.15154/z563-zd24). Briefly, the ABCD Study enrolled 11,880 children 9 to 10 years of age (mean age = 9.49; 48% female) in a 10-year longitudinal study at 21 different study sites across the United States.

### Participants

Children were recruited at 21 study sites from elementary schools (private, public, and charter schools) across the United States in a sampling design that aimed to represent the nationwide sociodemographic diversity (Garavan et al., 2018). All experimental and consent procedures were approved by the institutional review board and human research protections programs at the University of California San Diego. Each participant provided written assent, and their legal guardian provided written consent for them to participate in the study. Here, we use a subset of data from the ABCD Study, including magnetic resonance imaging (MRI), participants’ sex at birth, socio-demographics, and pubertal development, and sex hormones. Data include assessments from two time points: baseline enrollment and year 2 follow-up. Measures of participants’ sex, demographics from the initial study visit (i.e., at ages 9-11 years), while pubertal development, salivary sex hormones and MRI data were taken from both baseline and 2-year follow-up (i.e., at ages 11-13 years). Exclusion criteria for the ABCD Study included lack of English proficiency, severe sensory, neurological, medical or intellectual limitations, and inability to complete an MRI scan (Casey et al., 2018; Garavan et al., 2018). For this study, we further excluded participants who: had incidental neurological findings evident in their scans, failed the ABCD imaging or FreeSurfer quality control procedures (Hagler et al., 2019). Only images without clinically significant incidental findings, that passed all ABCD quality-control parameters, with low head motion (mean framewise displacement < 2mm) were included in the following analyses. Participants from site 15 were excluded from analyses due to a scanner hardware error (https://doi.org/10.5281/zenodo.22134953). Participant inclusion flow diagrams are presented in Supplementary Figures 1 and 2. The sample characteristics for the current study are described in Table 1.

**Table 1.**
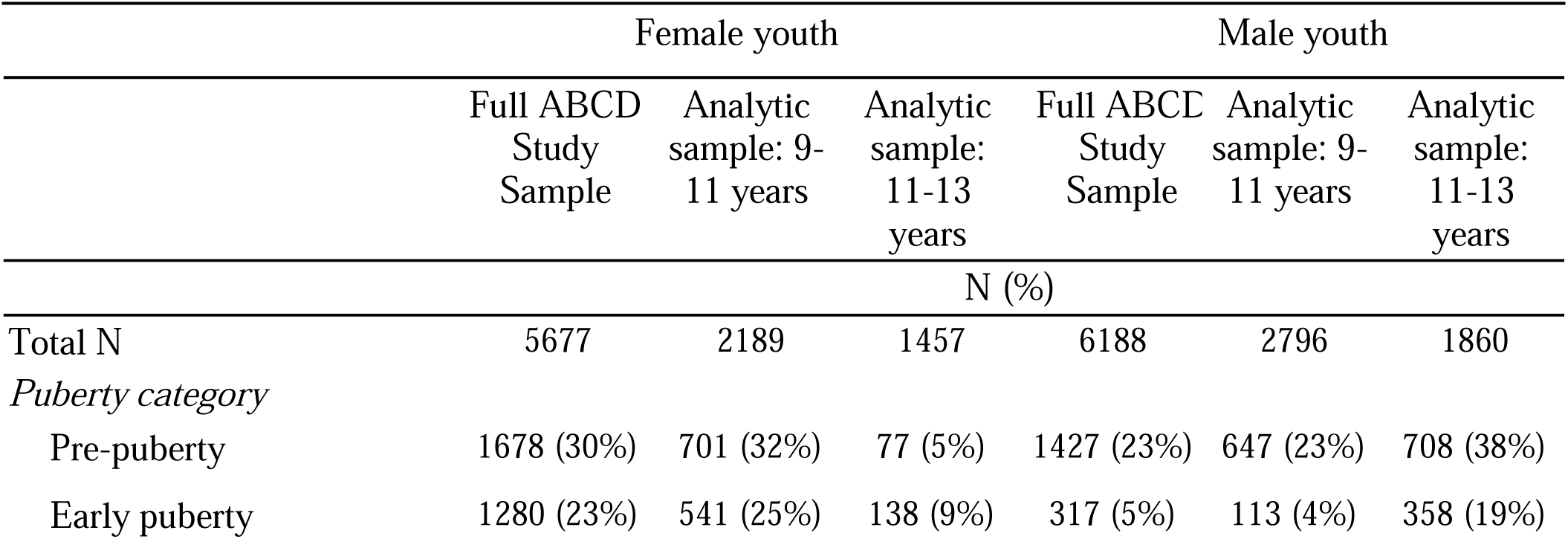
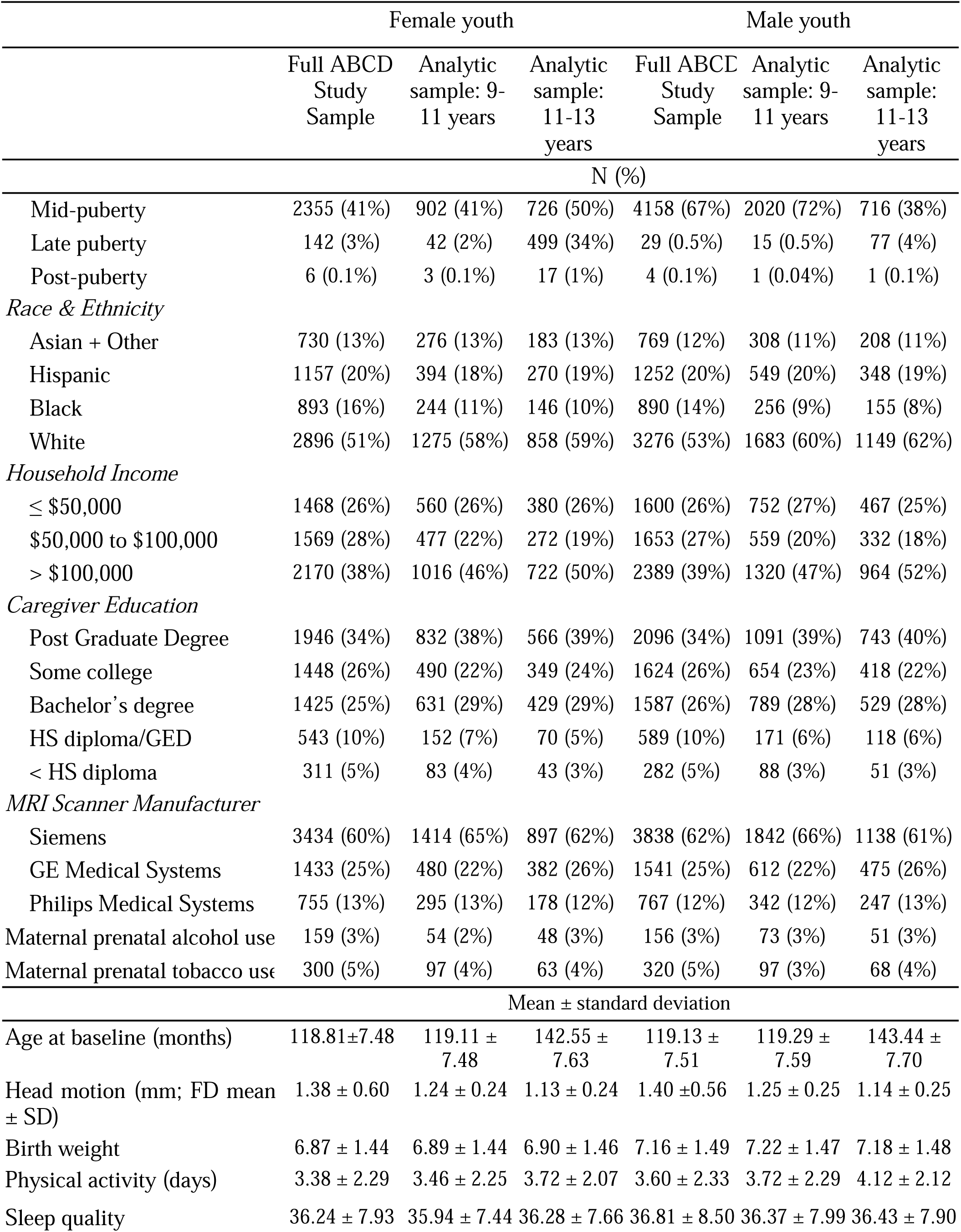
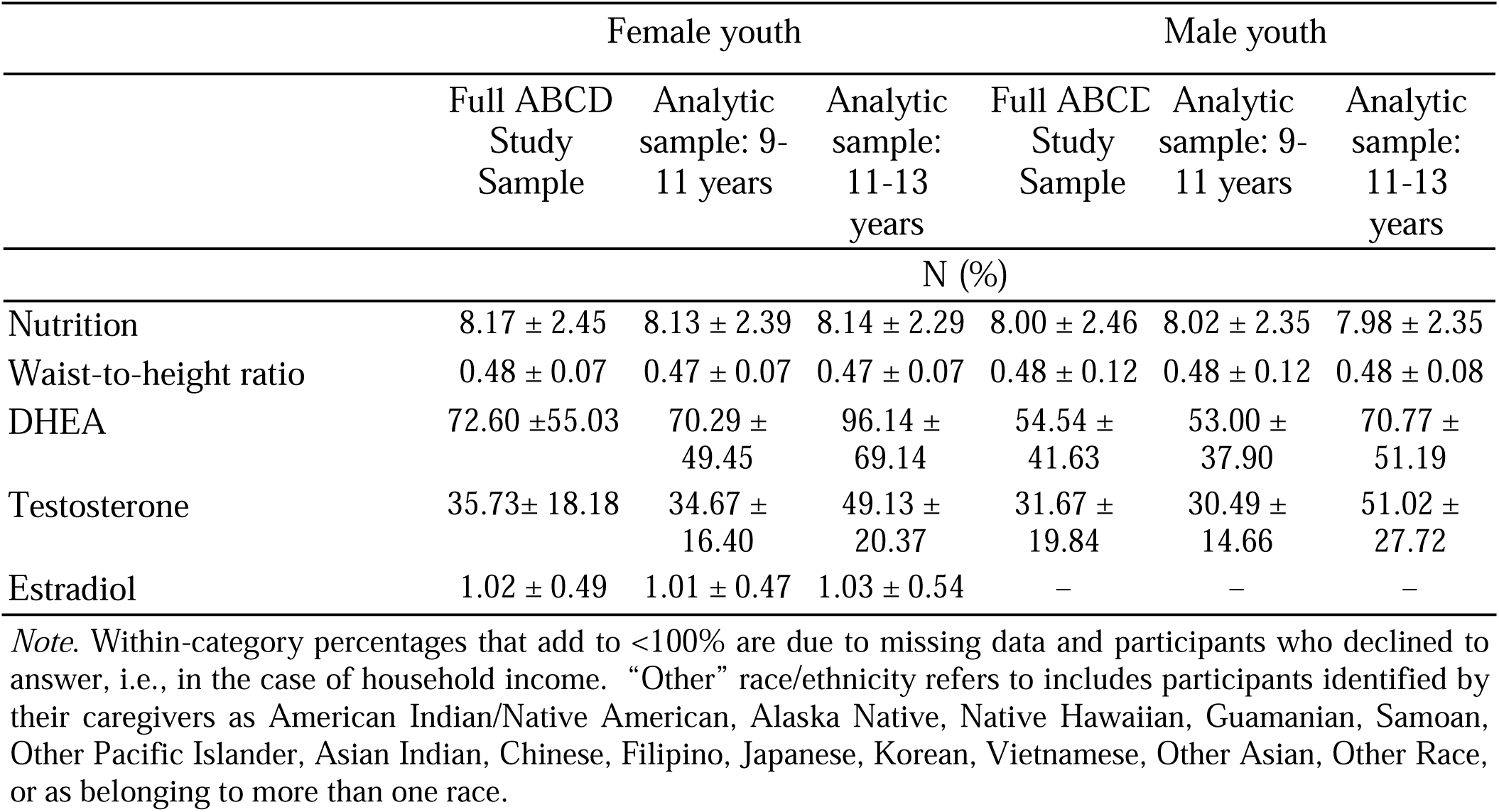
Sample Demographics.

|  | Female youth |  |  | Male youth |  |  |
|---|---|---|---|---|---|---|
|  | Full ABCD Study Sample | Analytic sample: 9-11 years | Analytic sample: 11-13 years | Full ABCD Study Sample | Analytic sample: 9-11 years | Analytic sample: 11-13 years |
|  | N (%) |  |  |  |  |  |
| Total N | 5677 | 2189 | 1457 | 6188 | 2796 | 1860 |
| *Puberty category* |  |  |  |  |  |  |
| Pre-puberty | 1678 (30%) | 701 (32%) | 77 (5%) | 1427 (23%) | 647 (23%) | 708 (38%) |
| Early puberty | 1280 (23%) | 541 (25%) | 138 (9%) | 317 (5%) | 113 (4%) | 358 (19%) |
|  | Female youth |  |  | Male youth |  |  |
|  | Full ABCD Study Sample | Analytic sample: 9-11 years | Analytic sample: 11-13 years | Full ABCD Study Sample | Analytic sample: 9-11 years | Analytic sample: 11-13 years |
|  | N (%) |  |  |  |  |  |
| Mid-puberty | 2355 (41%) | 902 (41%) | 726 (50%) | 4158 (67%) | 2020 (72%) | 716 (38%) |
| Late puberty | 142 (3%) | 42 (2%) | 499 (34%) | 29 (0.5%) | 15 (0.5%) | 77 (4%) |
| Post-puberty | 6 (0.1%) | 3 (0.1%) | 17 (1%) | 4 (0.1%) | 1 (0.04%) | 1 (0.1%) |
| *Race & Ethnicity* |  |  |  |  |  |  |
| Asian + Other | 730 (13%) | 276 (13%) | 183 (13%) | 769 (12%) | 308 (11%) | 208 (11%) |
| Hispanic | 1157 (20%) | 394 (18%) | 270 (19%) | 1252 (20%) | 549 (20%) | 348 (19%) |
| Black | 893 (16%) | 244 (11%) | 146 (10%) | 890 (14%) | 256 (9%) | 155 (8%) |
| White | 2896 (51%) | 1275 (58%) | 858 (59%) | 3276 (53%) | 1683 (60%) | 1149 (62%) |
| *Household Income* |  |  |  |  |  |  |
| ≤ \$50,000 | 1468 (26%) | 560 (26%) | 380 (26%) | 1600 (26%) | 752 (27%) | 467 (25%) |
| $50,000 to $100,000 | 1569 (28%) | 477 (22%) | 272 (19%) | 1653 (27%) | 559 (20%) | 332 (18%) |
| > \$100,000 | 2170 (38%) | 1016 (46%) | 722 (50%) | 2389 (39%) | 1320 (47%) | 964 (52%) |
| *Caregiver Education* |  |  |  |  |  |  |
| Post Graduate Degree | 1946 (34%) | 832 (38%) | 566 (39%) | 2096 (34%) | 1091 (39%) | 743 (40%) |
| Some college | 1448 (26%) | 490 (22%) | 349 (24%) | 1624 (26%) | 654 (23%) | 418 (22%) |
| Bachelor’s degree | 1425 (25%) | 631 (29%) | 429 (29%) | 1587 (26%) | 789 (28%) | 529 (28%) |
| HS diploma/GED | 543 (10%) | 152 (7%) | 70 (5%) | 589 (10%) | 171 (6%) | 118 (6%) |
| < HS diploma | 311 (5%) | 83 (4%) | 43 (3%) | 282 (5%) | 88 (3%) | 51 (3%) |
| *MRI Scanner Manufacturer* |  |  |  |  |  |  |
| Siemens | 3434 (60%) | 1414 (65%) | 897 (62%) | 3838 (62%) | 1842 (66%) | 1138 (61%) |
| GE Medical Systems | 1433 (25%) | 480 (22%) | 382 (26%) | 1541 (25%) | 612 (22%) | 475 (26%) |
| Philips Medical Systems | 755 (13%) | 295 (13%) | 178 (12%) | 767 (12%) | 342 (12%) | 247 (13%) |
| Maternal prenatal alcohol use | 159 (3%) | 54 (2%) | 48 (3%) | 156 (3%) | 73 (3%) | 51 (3%) |
| Maternal prenatal tobacco use | 300 (5%) | 97 (4%) | 63 (4%) | 320 (5%) | 97 (3%) | 68 (4%) |
|  | Mean ± standard deviation |  |  |  |  |  |
| Age at baseline (months) | 118.81±7.48 | 119.11 ± 7.48 | 142.55 ± 7.63 | 119.13 ± 7.51 | 119.29 ± 7.59 | 143.44 ± 7.70 |
| Head motion (mm; FD mean ± SD) | 1.38 ± 0.60 | 1.24 ± 0.24 | 1.13 ± 0.24 | 1.40 ±0.56 | 1.25 ± 0.25 | 1.14 ± 0.25 |
| Birth weight | 6.87 ± 1.44 | 6.89 ± 1.44 | 6.90 ± 1.46 | 7.16 ± 1.49 | 7.22 ± 1.47 | 7.18 ± 1.48 |
| Physical activity (days) | 3.38 ± 2.29 | 3.46 ± 2.25 | 3.72 ± 2.07 | 3.60 ± 2.33 | 3.72 ± 2.29 | 4.12 ± 2.12 |
| Sleep quality | 36.24 ± 7.93 | 35.94 ± 7.44 | 36.28 ± 7.66 | 36.81 ± 8.50 | 36.37 ± 7.99 | 36.43 ± 7.90 |
|  | Female youth |  |  | Male youth |  |  |
|  | Full ABCD Study Sample | Analytic sample: 9-11 years | Analytic sample: 11-13 years | Full ABCD Study Sample | Analytic sample: 9-11 years | Analytic sample: 11-13 years |
|  | N (%) |  |  |  |  |  |
| Nutrition | 8.17 ± 2.45 | 8.13 ± 2.39 | 8.14 ± 2.29 | 8.00 ± 2.46 | 8.02 ± 2.35 | 7.98 ± 2.35 |
| Waist-to-height ratio | 0.48 ± 0.07 | 0.47 ± 0.07 | 0.47 ± 0.07 | 0.48 ± 0.12 | 0.48 ± 0.12 | 0.48 ± 0.08 |
| DHEA | 72.60 ±55.03 | 70.29 ± 49.45 | 96.14 ± 69.14 | 54.54 ± 41.63 | 53.00 ± 37.90 | 70.77 ± 51.19 |
| Testosterone | 35.73± 18.18 | 34.67 ± 16.40 | 49.13 ± 20.37 | 31.67 ± 19.84 | 30.49 ± 14.66 | 51.02 ± 27.72 |
| Estradiol | 1.02 ± 0.49 | 1.01 ± 0.47 | 1.03 ± 0.54 | – | – | – |

*Note*. Within-category percentages that add to <100% are due to missing data and participants who declined to answer, i.e., in the case of household income. “Other” race/ethnicity refers to includes participants identified by their caregivers as American Indian/Native American, Alaska Native, Native Hawaiian, Guamanian, Samoan, Other Pacific Islander, Asian Indian, Chinese, Filipino, Japanese, Korean, Vietnamese, Other Asian, Other Race, or as belonging to more than one race.

### Neuroimaging Data

A harmonized data protocol was utilized across sites with either a Siemens, Phillips, or GE 3T MRI scanner. Motion compliance training, as well as real-time, prospective motion correction was used to reduce motion distortion.

#### Structural MRI: Acquisition, Processing, and Quality Control

T1-weighted (T1w) images were acquired using a magnetization-prepared rapid acquisition gradient echo (MPRAGE) sequence with 176 slices and 1mm^3^ isotropic resolution (Casey et al., 2018). FreeSurfer (v7.1.1) was used to preprocess T1w images (e.g., skull stripping, tissue segmentation, surface reconstruction) and apply cortical and subcortical parcellations (Dale et al., 1999; Destrieux et al., 2010; Fischl et al., 1999, 2001; Segonne et al., 2007; Ségonne et al., 2004). Additional details are published elsewhere (Hagler et al., 2019).

#### Diffusion Weighted Imaging (DWI): Acquisition, Preprocessing, and Quality Control

Prior to each DWI acquisition, a fieldmap was acquired to account for B0 distortion correction (Casey et al., 2018). Multi-shell diffusion-weighted imaging was acquired using an echo planar imaging (EPI) sequence (multiband factor = 3) with 1.7mm^3^ isotropic voxels (Moeller et al., 2010; Setsompop et al., 2012), seven b=0 frames, and 96 diffusion directions at 4 b-values (6 at b = 500 s/mm^2^, 15 at b = 1000 s/mm^2^, 15 at b = 2000 s/mm^2^, and 60 at b = 3000 s/mm^2^). Image processing included eddy current distortion correction (Zhuang et al., 2006), head motion correction (Hagler et al., 2019), diffusion gradient adjustments for head motion (Hagler et al., 2009; Leemans & Jones, 2009), tensor estimation to identify and replace head motion outlier frames (Chang et al., 2005), B0 distortion correction (Andersson et al., 2003; S. M. Smith et al., 2004), gradient nonlinearity distortion correction (Jovicich et al., 2006), coregistration to T1w images (Wells et al., 1996), and resampling to a standard orientation with 1.7mm isotropic voxels.

#### Restricted Spectrum Imaging (RSI)

Restricted spectrum imaging (RSI) is an advanced modeling technique that utilizes all 96 directions collected as part of ABCD’s multi-shell acquisition protocol. RSI provides detailed information regarding both the extracellular and intracellular compartments of neural white matter (White et al., 2012, 2013, 2014). To evaluate cellular microarchitecture, we utilized two RSI model outputs: restricted normalized isotropic signal fraction (RNI) and restricted normalized directional signal fraction (RND). RNI measures intracellular isotropic diffusion. A higher RNI value could indicate an increase in the number of support cells or swelling of support cells, such as activated astrocytes or microglia (Palmer et al., 2022). RND measures intracellular anisotropic diffusion and likely indicates diffusion within axons and dendrites. Higher RND values indicate more myelination and/or axonal packing (Palmer et al., 2022). Mean RSI values (ranging from 0 to 1) were calculated for 74 cortical gray matter regions per hemisphere (gyral and sulcal) defined by the Destrieux atlas (Destrieux et al., 2010), 13 subcortical gray matter regions (Fischl et al., 2002), and 35 white matter fiber tracts defined using AtlasTrack (Hagler et al., 2009) for a total of 196 regional microstructure features each for RNI and RND.

### Puberty & Hormone Estimates

The Pubertal Development Scale (PDS) was used to measure perceived pubertal development (Petersen et al., 1988) and was completed by caregivers about the participants and by youth about themselves (Barch et al., 2018; Cheng et al., 2021; Herting et al., 2021). Because 9-10 year old youth estimates were previously found to be unreliable (Herting et al., 2021), the present study used caregiver reports. The PDS comprises three questions asked about all youth–regarding height growth, the appearance of body hair, and skin changes–and two sex-specific questions. For female youth these address breast development and menarche; for male youth they address voice changes and facial hair. Each question is a Likert item from 1 (has not begun) to 4 (definitely begun). Scores used here represent the average across these five questions.

Dehydroepiandrosterone (DHEA), testosterone (T), and estradiol (E_2_) levels were measured from saliva samples collected during study visits via passive drool (Granger et al., 2012; Herting et al., 2021). Participants refrained from eating for at least 30 minutes before sample collection and rinsed their mouths with water at 10 minutes prior, to mitigate contamination. Samples were cooled after collection and either immediately frozen (i.e., in a-80C or-20C freezer) or refrigerated at 4C until neurocognitive testing was completed, and then frozen. Samples were analyzed by Salimetrics (Carlsbad, CA) between 2 and 6 months after collection. Procedures followed protocols for commercially available immunoassays, in duplicate, and assessed levels of DHEA (6.1 to 600 pg/mL with a lower limit of 1pg/mL) and T (10.2 to 1,000 pg/mL with a lower limit of 5pg/mL) in all samples and levels of E_2_ (1 to 32 pg/mL with a lower limit of 0.1pg/mL) only in female participants’ samples.

### Bias-Adjusting and Precision Variables

A minimally sufficient set of bias-adjusting variables was identified previously, using a directed acyclic graph that modeled potential biasing paths between pubertal development and MRI phenotypes (Supplementary Figure 3). This set included sociodemographic variables (i.e., youth race/ethnicity, average household income in USD, and highest household education), prenatal factors (i.e., maternal tobacco and alcohol use during pregnancy, birth weight, maternal age), and physical health (i.e., recent physical activity, nutrition, waist-to-height ratio, sleep quality), in addition to youth age. To this, we added precision variables to account for differences in salivary hormone estimates (i.e., time since midnight, collection duration, elapsed time before sample was frozen, recent physical activity, and recent caffeine intake) and to account for differences in MRI data acquisition (i.e., MRI scanner manufacturer, handedness, head motion during dMRI scan, data collection site). See Table 2 for details.

**Table 2.** Bias-adjusting and precision variables included in each regression model.

| Measure | Description | Reference |
|---|---|---|
| *Perinatal factors* |  |  |
| Maternal alcohol use | Once you/biological mother knew you/she were pregnant, were you/biological mother using… alcohol? |  |
| Maternal tobacco use | Once you/biological mother knew you/she were pregnant, were you/biological mother using… tobacco? |  |
| Maternal age | Caregiver reported, in years |  |
| Birth weight | Caregiver reported, in pounds |  |
| *Physical health* |  |  |
| Physical activity | Reported by youth as number of days in the preceding week with more than 60 minutes of physical activity, per Youth Risk Behavior Survey | (CDC, 2018) |
| Sleep quality | Sleep Disturbance Scale for Children | (Bruni et al., 1996) |
| Waist-to-height ratio | Waist circumference divided by height (each in inches, average of 2 measurements) | (Agbaje, 2024) |
| Nutrition | Child Nutrition Assessment | (Liu et al., 2021; Nagata et al., 2024) |
| *Sociodemographic factors* |  |  |
| Age | Number of months elapsed between youth date of birth and interview date | PhenX Toolkit (Hamilton et al., 2011) |
| Household income | 4-level categorical summary: >\$100K, \$50K-\$100K, ≤\$50K, or Don’t Know/Refuse to Answer |  |
| Caregiver education | 5-level categorical summary: Post-Graduate, Bachelor, Some College, High School Diploma/GED, or <High School Diploma |  |
| Youth race/ethnicity | 5-level categorical summary: non-Hispanic white, non-Hispanic Black, Hispanic, non-Hispanic Asian/Other |  |
| *Precision variables – Saliva* |  |  |
| Collection time | Minutes since midnight | (Herting et al., 2021) |
| Collection duration | Minutes from beginning to end of active sample collection |  |
| Elapsed time before sample was frozen | Minutes from sample collection end time to storage in on-site freezer |  |
| Recent caffeine intake | Caffeine consumption in the 12 hours preceding sample collection (yes/no) |  |
| Recent physical activity | Vigorous physical activity in the 12 hours preceding sample collection (yes/no) |  |
| *Precision variables – MRI* |  |  |
| MRI scanner manufacturer | 3-level categorical variable: General Electric, Philips, or SIEMENS |  |
| Handedness | Edinburgh Handedness Inventory summary |  |
| Head motion during dMRI scan | Average framewise displacement, in mm | (Power et al., 2012; Satterthwaite et al., 2013) |
| Data collection site | 20-level categorical variable,* indicating site at which data were acquired |  |

*Note.* Data from only 20 ABCD Study sites was used due to a scanner hardware malfunction at site 15 (https://doi.org/10.5281/zenodo.22134953).

### Analyses

Prior to data analysis, an analysis plan was pre-registered on the Open Science Framework (OSF; osf.io/fjcrw/). Code used to perform the following analyses is available on Github (github.com/BottenLab/microPuberty) (Bottenhorn, 2026). Outlier participants were identified using unsupervised multivariate anomaly detection, by comparing the local density (i.e., distance of each participant to their 10 nearest neighbors, the 10 most similar participants based on microarchitecture and hormone data) of each participant to the local densities of their neighbors. Participants with lower densities than their neighbors (i.e., more dissimilar to other participants) were considered outliers and removed from subsequent analyses.

Individual differences in intracellular diffusion (i.e., RNI, RND) related to pubertal hormone levels were identified using multi-task elastic net regression (i.e., linear regression with *L*_1_ and *L*_2_ penalties) in scikit-learn (v1.0.1) (Pedregosa et al., 2011). Briefly, this approach estimates coefficients via gradient descent for multiple regression problems simultaneously, returning a sparse set of predictors that are related to all outcomes (though their coefficients may vary by outcome; Figure 2a). Here, predictors include intracellular diffusion values per brain region/tract, in addition to the bias-adjusting and precision variables noted above, with separate models fit per phenotype (i.e., RNI, RND). Multi-task regression outcomes were both average PDS scores and hormone levels with separate models fit per hormone (i.e., DHEA, T, E_2_). Data were split into training and testing subsets in an outer cross-validation loop by study site, such that the training data included participants from 17 of 20 sites and the testing data included participants from the remaining three sites (Figure 2b). Models were trained with built-in hyperparameter tuning with 5-fold cross-validation (i.e., inner CV loop) to identify the optimal *L*_1_-to-*L*_2_ ratio. Trained models were then fit to testing data and model performance was estimated as the coefficient of determination (*R*^2^). Significance testing was not performed for models or individual coefficients, as the elastic net penalty intentionally shrinks estimates and violates the unbiasedness assumptions of inferential ordinary least squares regression statistics. Instead of reporting model significance as *p*-values, we evaluated the explanatory power of each joint-outcome multi-task model using out-of-sample variance explained across iterative cross-validation folds. This provides a more interpretable, robust measure of generalized predictive performance. Finally, the relevance of each predictor to pubertal hormone levels was estimated using feature permutation testing in the testing data (Figure 2c), which shuffled values of each predictor, in turn, and calculated the resulting *R*^2^. Values were randomly shuffled 50 times for each predictor, within 1000 iterations of the outer CV loop, for a total of 50,000 estimates of changes in performance due to shuffled values (i.e., *relevance*) per predictor. Regional relevance was calculated as the proportion of total model performance (i.e., *R*^2^) attributable to randomizing microarchitecture values. Hyperparameter results and overall model performance are displayed in Table 4.

**Figure 2.**
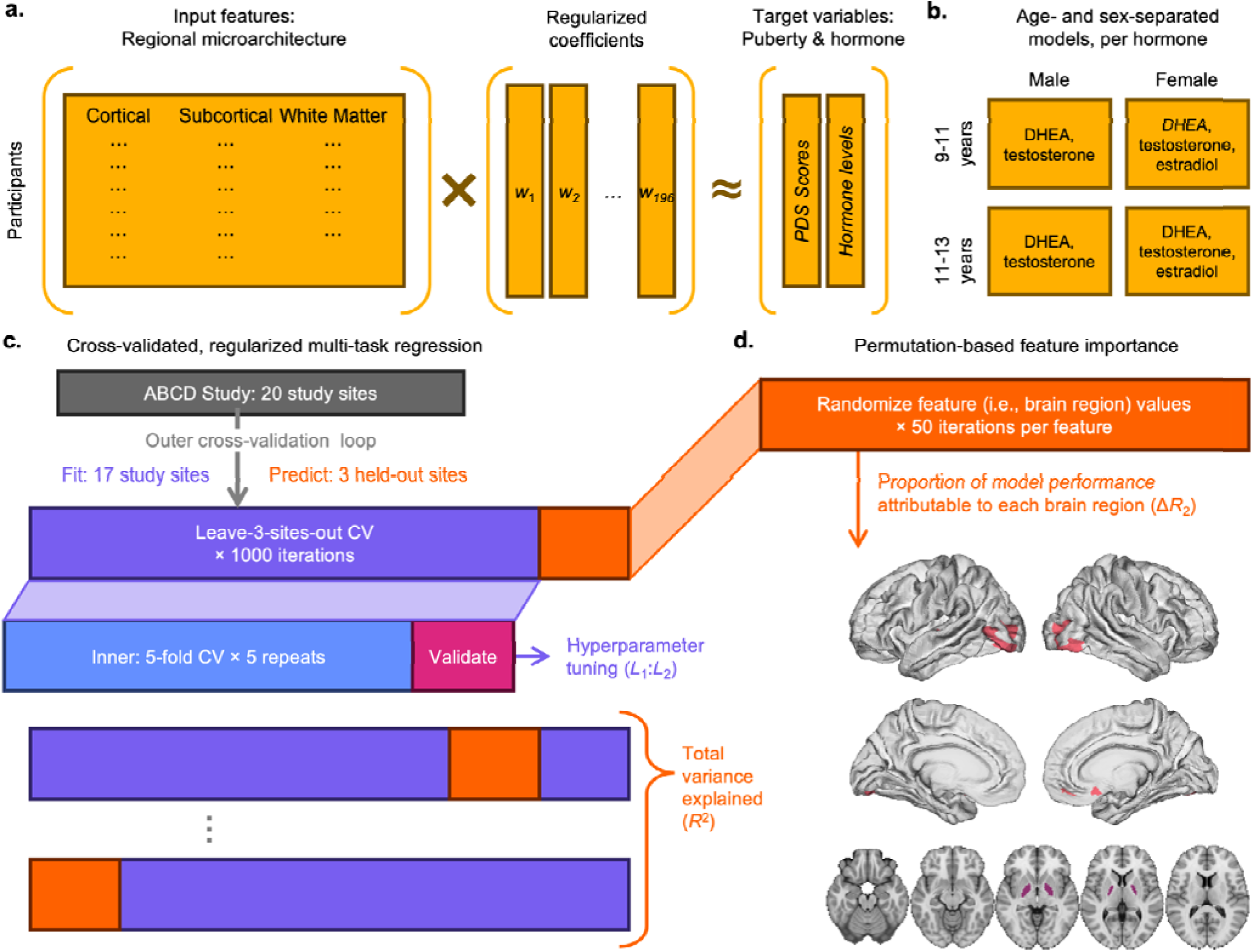
Conceptual diagram of multi-task regression pipeline: Predicting pubertal hormones from brain microarchitecture. (A) Here, we used regularized multi-task regression with regional cortical, subcortical, and white matter micro-architecture (i.e., cellular density, RNI; neurite density, RND) as input features, with both PDS scores and hormone levels (i.e., DHEA, testosterone, estradiol) as target variables. (B) Age-and sex-stratified models were run regressing PDS scores and levels of each hormone, in turn, on brain microarchitecture for a total of 10 models. (C) Regression models were trained in a nested cross-validation (CV) framework. A leave-3-sites-out outer CV loop fit regression models on data from 17 sites–using an inner 5-fold CV loop to tune shrinkage ratio (*L*_1_:*L*_2_)– and predicted target variables in 3 sites, across 1000 iterations. Model performance was calculated in test data as the proportion of variance in PDS scores and hormone levels by the full model (*R*^2^). (D) Target variable predictions were repeated 50 times per feature, with randomized feature variables in each permutation, to calculate each feature’s relative importance to model predictions as the change in variance explained (Δ*R*^2^).

## Results

At 9-11 years of age, dMRI from *N* = 4,355 female youth and *N* = 3,129 male youth passed image quality control. At 11-13 years of age, dMRI data from *N* = 3,129 female youth and *N* = 3,574 male youth passed image quality control. After removing participants who were multivariate outliers on neuroendocrine variables (i.e., dMRI, hormones, PDS) and participants with incomplete data on model variables (i.e., neuroendocrine confounders, precision variables), the final analytic samples included *N* = 2,189 female youth at 9-10 years of age, *N* = 1,457 female youth at 11-13 years of age, *N* = 2,796 male youth at 9-11 years of age, and *N* = 1,860 male youth at 11-13 years of age. Proportions of missing data across the sample for all model variables are provided in Supplementary Table 1. Briefly, proportions of missing data were below 5 percent for most variables. Only birth weight (∼10%), dMRI data (6% at 9-11 years of age, 26% at 11-13 years of age), nutrition scores (14-16%), and hormone estimates (DHEA: 12% at 9-11 years of age, 36% at 11-13 years of age; T: 12% at 9-11 years of age, 35% at 11-13 years of age) had greater than 5% of values missing. We also calculated product-moment correlations between PDS scores and salivary levels of each hormone for each male and female samples at each age range (Table 3).

**Table 3.** Outcome (i.e., PDS, hormones) descriptive statistics and correlations across age and sex.

|  | Male 9-11 years | Male, 11-13 years | Female, 9-11 years | Female, 11-13 years |
|---|---|---|---|---|
| PDS Average | 1.4 [1.2, 1.6] | 1.6 [1.4, 2.05] | 1.6 [1.4, 2.0] | 2.4 [2.0, 3.2] |
| DHEA | 43.65 [25.27, 70.38] | 58.39 [33.67, 92.89] | 58.12 [34.1, 94.36] | 80.06 [47.81, 127.79] |
| Testosterone | 28.02 [19.84, 38.75] | 44.24 [31.58, 64.08] | 32.19 [22.41, 43.29] | 46.71 [34.15, 60.55] |
| Estradiol | – | – | 0.97 [0.68, 1.3] | 0.98 [0.65, 1.33] |
| *Correlation with PDS average* |  |  |  |  |
| DHEA | 0.161, 1.02×10<sup>-17</sup> | 0.297, 3.44×10<sup>-39</sup> | 0.301, 6.01×10<sup>-47</sup> | 0.344, 1.05×10<sup>-41</sup> |
| Testosterone | 0.143, 2.52×10<sup>-14</sup> | 0.463, 1.14×10<sup>-99</sup> | 0.281, 5.61×10<sup>-41</sup> | 0.313, 1.57×10<sup>-34</sup> |
| Estradiol | – | – | 0.098, 4.37×10<sup>-06</sup> | 0.24, 1.48×10<sup>-20</sup> |

*Note*. Hormones were measured in pg/mL. Descriptive statistics follow the form “median [25%ile, 75%ile]”; correlations of the form “*r<sub>S</sub>*, *p*(*r<sub>S</sub>*)”. All correlations are significant at *α* < 0.01.

**Table 4.**
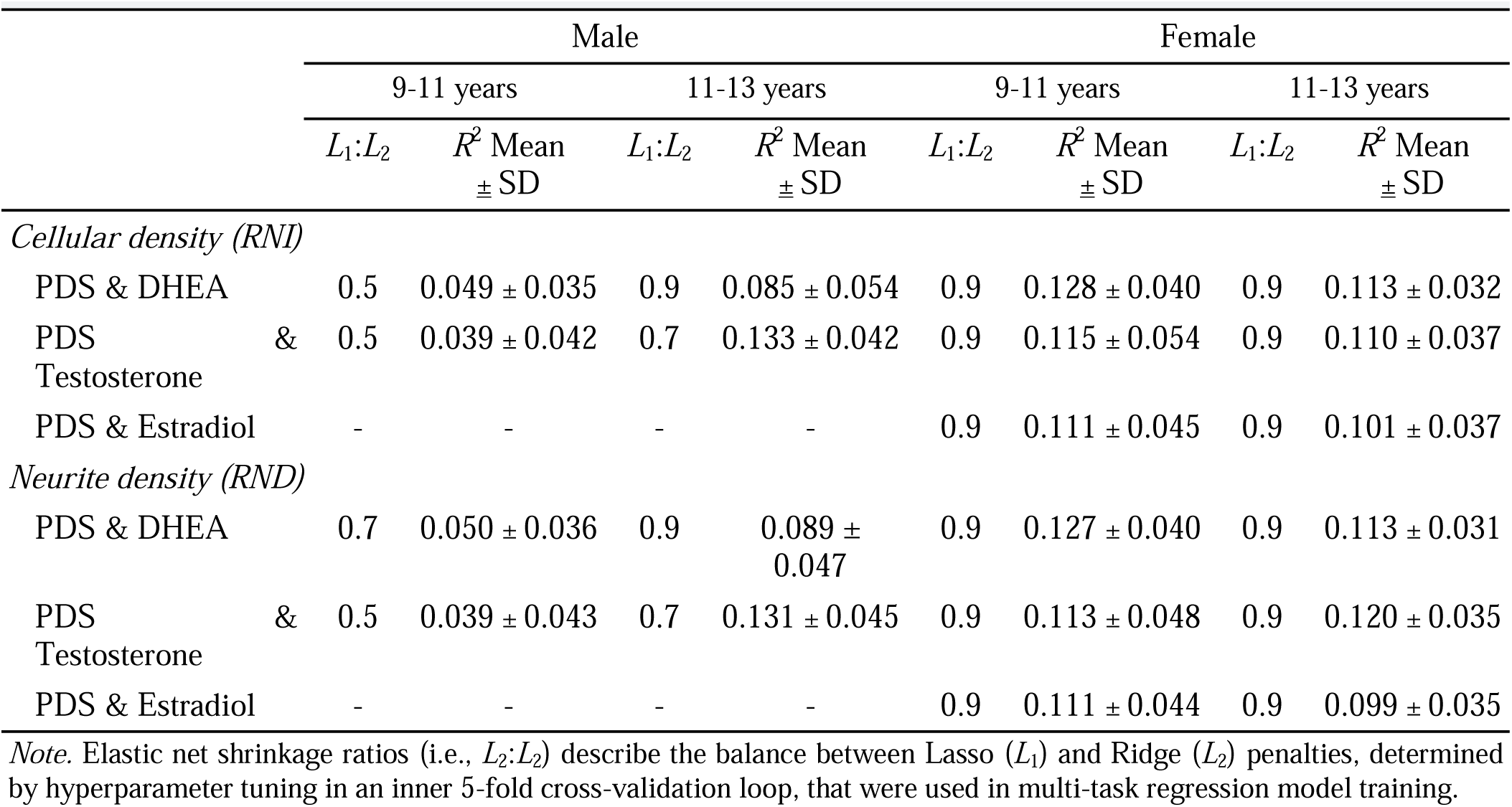
Tuned parameters and variance explained for RNI and RND models.

|  | Male |  |  |  | Female |  |  |  |
|---|---|---|---|---|---|---|---|---|
|  | 9-11 years |  | 11-13 years |  | 9-11 years |  | 11-13 years |  |
|  | *L*<sub>1</sub>:*L*<sub>2</sub> | *R*<sup>2</sup> Mean ± SD | *L*<sub>1</sub>:*L*<sub>2</sub> | *R*<sup>2</sup> Mean ± SD | *L*<sub>1</sub>:*L*<sub>2</sub> | *R*<sup>2</sup> Mean ± SD | *L*<sub>1</sub>:*L*<sub>2</sub> | *R*<sup>2</sup> Mean ± SD |
| *Cellular density (RNI)* |  |  |  |  |  |  |  |  |
| PDS & DHEA | 0.5 | 0.049 ± 0.035 | 0.9 | 0.085 ± 0.054 | 0.9 | 0.128 ± 0.040 | 0.9 | 0.113 ± 0.032 |
| PDS & Testosterone | 0.5 | 0.039 ± 0.042 | 0.7 | 0.133 ± 0.042 | 0.9 | 0.115 ± 0.054 | 0.9 | 0.110 ± 0.037 |
| PDS & Estradiol | - | - | - | - | 0.9 | 0.111 ± 0.045 | 0.9 | 0.101 ± 0.037 |
| *Neurite density (RND)* |  |  |  |  |  |  |  |  |
| PDS & DHEA | 0.7 | 0.050 ± 0.036 | 0.9 | 0.089 ± 0.047 | 0.9 | 0.127 ± 0.040 | 0.9 | 0.113 ± 0.031 |
| PDS & Testosterone | 0.5 | 0.039 ± 0.043 | 0.7 | 0.131 ± 0.045 | 0.9 | 0.113 ± 0.048 | 0.9 | 0.120 ± 0.035 |
| PDS & Estradiol | - | - | - | - | 0.9 | 0.111 ± 0.044 | 0.9 | 0.099 ± 0.035 |

*Note.* Elastic net shrinkage ratios (i.e., *L*<sub>2</sub>:*L*<sub>2</sub>) describe the balance between Lasso (*L*<sub>1</sub>) and Ridge (*L*<sub>2</sub>) penalties, determined by hyperparameter tuning in an inner 5-fold cross-validation loop, that were used in multi-task regression model training.

Across hormones and microarchitecture measures, inner-CV loop hyperparameter tuning determined that regression models in 9–11-year-old male youth had consistently lower optimal shrinkage ratios (*L*_1_:*L*_2_ = [0.5-0.7]) than did 11-13-year-old male youth (*L*_1_:*L*_2_ = [0.7–0.9]) or female youth at each age range ((*L*_1_:*L*_2_ = 0.9); Table 4). This means that regression models in younger male youth will be less sparse than regression models in older male youth or in female youth.

A higher shrinkage ratio indicates a sparser model (i.e., more feature coefficients driven to zero). The coefficient of determination (i.e., *R*^2^) was calculated when trained models were applied to testing data in each outer CV loop and describes how much of the variability in PDS scores and hormone levels is predictable by microarchitecture, neuroendocrine confounds, and precision variables.

For each multi-task regression (i.e., age-and sex-stratified regression of RNI and RND per hormone), we report only brain regions that were relevant to PDS and hormone predictions in over 85% of leave-3-sites-out cross-validation model iterations (Supplementary Tables 1-5).

### Microarchitecture related to pubertal DHEA differences

#### Cellular density (RNI)

For male youth aged 9-11 years (Figure 3a), individual differences in RNI accounted for 4.9% of the model-explained variance in pubertal DHEA levels (*R*^2^ = 0.049 + 0.035). Three of these regions were relevant in all model iterations: the bilateral middle occipital sulcus and lunatus sulcus and the right medial orbital sulcus. At ages 11-13 years (Figure 3b), individual differences in RNI accounted for 8.5% of model-explained pubertal DHEA variance (*R*^2^ = 0.085 ± 0.054). The right pallidum was relevant in 97% of model iterations, while the left pallidum and the left subcallosal gyrus were relevant in every model iteration. RNI differences in these regions accounted for 9.1-9.2% of model-explained pubertal DHEA variance.

**Figure 3.**
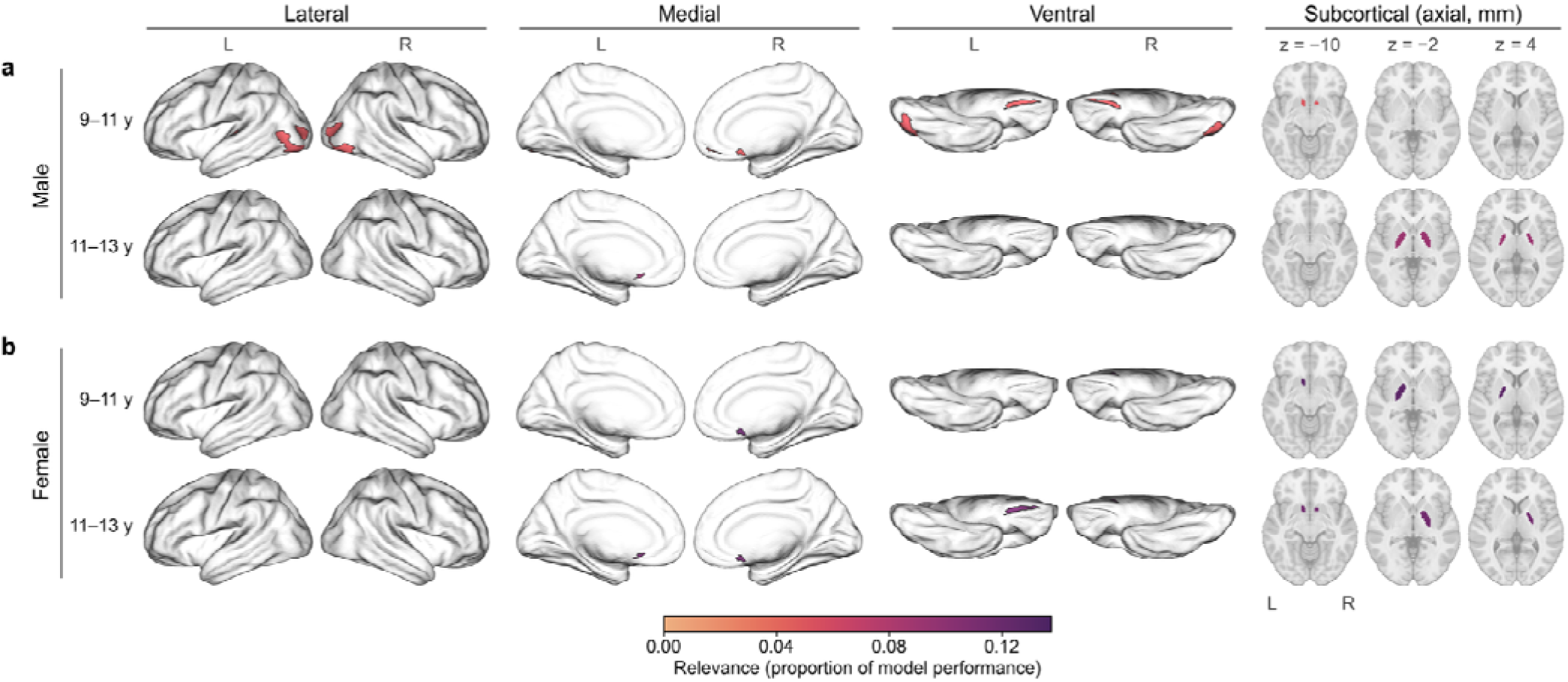
Regions exhibiting individual differences in RNI that were consistently relevant to pubertal DHEA predictions across model iterations. Color-coding indicates the average change in model performance associated with randomizing participant values for each region, across 50 permutations (per region, per cross-validation iteration.

For female youth aged 9-11 years (Figure 3b), individual differences in RNI accounted for 12.8% of the model-explained variance in pubertal DHEA levels (*R*^2^ = 0.128 ± 0.040). The right nucleus accumbens was relevant in every model iteration, and the left pallidum and subcallosal gyrus were relevant in 98% of model iterations. Individual differences in RNI of these regions accounted for 12.7% of model-explained pubertal DHEA variance. At 11-13 years (Figure 3b), individual differences in RNI accounted for 11.3% of the model-explained variance in pubertal DHEA levels (*R*^2^ = 0.113 ± 0.032). Bilateral nucleus accumbens and subcallosal gyri were relevant in all model iterations, while the right pallidum and left medial orbital sulcus were relevant in ≤95% of model iterations. Individual differences in RNI of these regions accounted for 11.3-11.4% of model-explained variance in pubertal DHEA.

At ages 9-11 years, differences in RNI of the left accumbens area were relevant to pubertal DHEA levels in both male and female youth. At ages 11-13 years, the right pallidum and left subcallosal gyrus were relevant in both sexes. Relevant regions from each model are presented in Figure 3 and Supplementary Table 1.

#### Neurite density (RND)

In male youth aged 9-11 years, individual differences in RND explained 5% of the variance in pubertal DHEA levels (*R*^2^ = 0.050 ± 0.036). Nine regions were relevant in all model iterations: the right inferior occipital gyrus and sulcus, left inferior part of the precentral sulcus, left middle frontal sulcus, right orbital sulci, bilateral planum temporale, left transverse temporal sulcus, left inferior occipital gyrus and sulcus, and right pallidum. Individual differences in the RND of these regions accounted for 4.9-5.4% of model-explained variance in pubertal DHEA levels. At ages 11-13 years, individual differences in RND explained 8.9% of the variance in pubertal DHEA levels (*R*^2^ = 0.089 ± 0.047), but no individual regions were consistently relevant.

For female youth aged 9-11 years, individual differences in RND explained 12.7% of the variance in pubertal DHEA levels (*R*^2^ = 0.127 ± 0.04). The left transverse temporal sulcus was relevant to pubertal DHEA predictions in 95% of model iterations and the pericallosal sulcus was relevant in 85% of iterations. Individual differences in the RND of these regions accounted for 12.7-12.8% of model-explained variance in pubertal DHEA levels. At ages 11-13 years, individual differences in RND explained 11.3% of the variance in pubertal DHEA levels (*R*^2^ = 0.113 ± 0.031). The right middle occipital sulcus and lunatus sulcus and the left cingulate cingulum were relevant in 91% and 88% of model iterations, respectively, and individual differences in these regions’ RND accounted for 10.9-11.4% of model-explained variance in pubertal DHEA.

At ages 9-11 years, the left pericallosal sulcus and the left transverse temporal sulcus were relevant for both male and female youth, but no regions were relevant for both sexes at ages 11-13 years. Relevant regions from each model are presented in Figure 4 and Supplementary Table 2.

**Figure 4.**
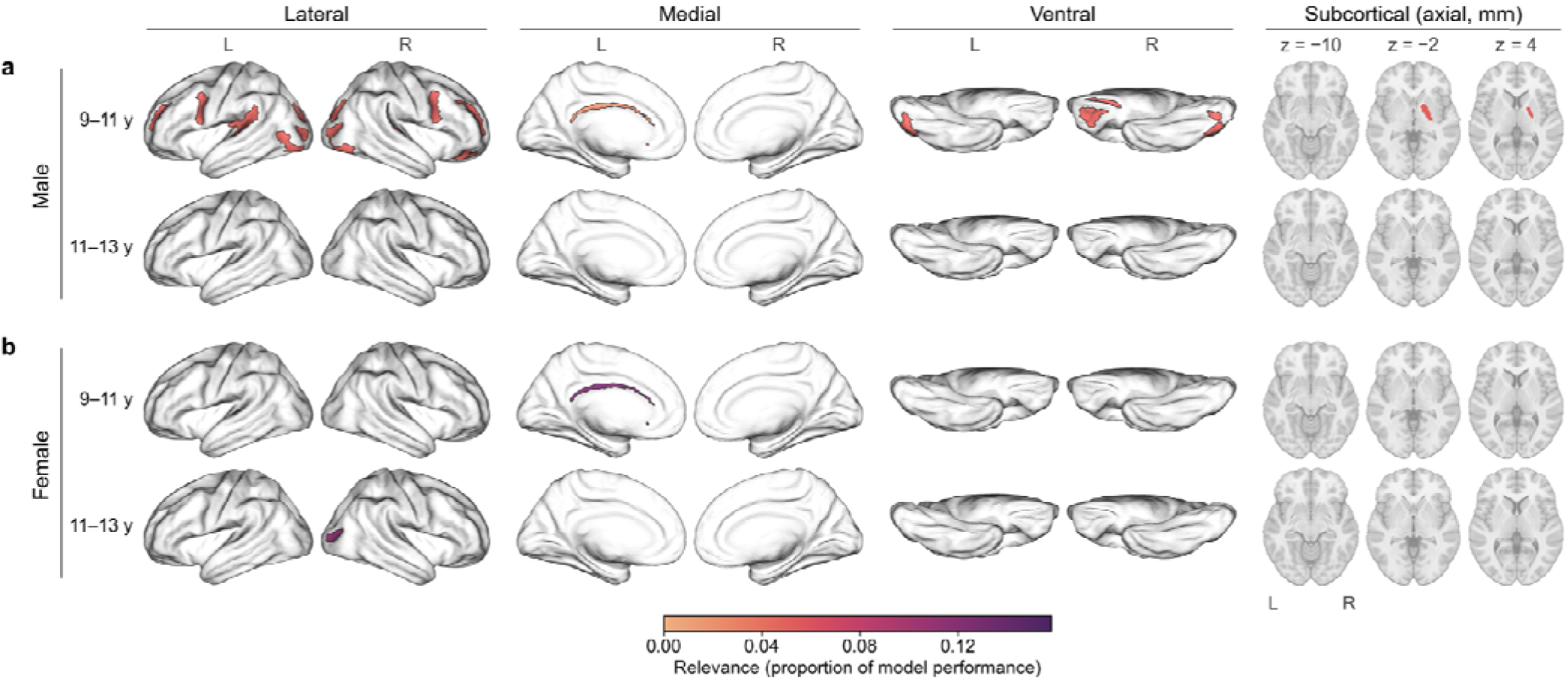
Regions exhibiting individual differences in RND that were consistently relevant to pubertal DHEA predictions across model iterations. Color-coding indicates the average change in model performance associated with randomizing participant values for each region, across 50 permutations (per region, per cross-validation iteration).

### Microarchitecture related to pubertal testosterone differences

#### Cellular density (RNI)

For male youth aged 9-11 years (Figure 5a), individual differences in RNI explained 3.9% of the variance in pubertal DHEA levels (R^2^= 0.039 ± 0.042). The left middle occipital sulcus and lunatus sulcus was relevant in 99% of model iterations, accounting for 4.0-4.3% of model-explained variance in pubertal testosterone. At ages 11-13 years, individual differences in RNI explained 13.3% of the variance in pubertal testosterone levels (R^2^ = 0.133 ± 0.042). The left subcallosal gyrus and left pallidum were relevant in ≥97% of model iterations and accounted for 13.6-13.7% of model-explained variance in pubertal testosterone.

**Figure 5.**
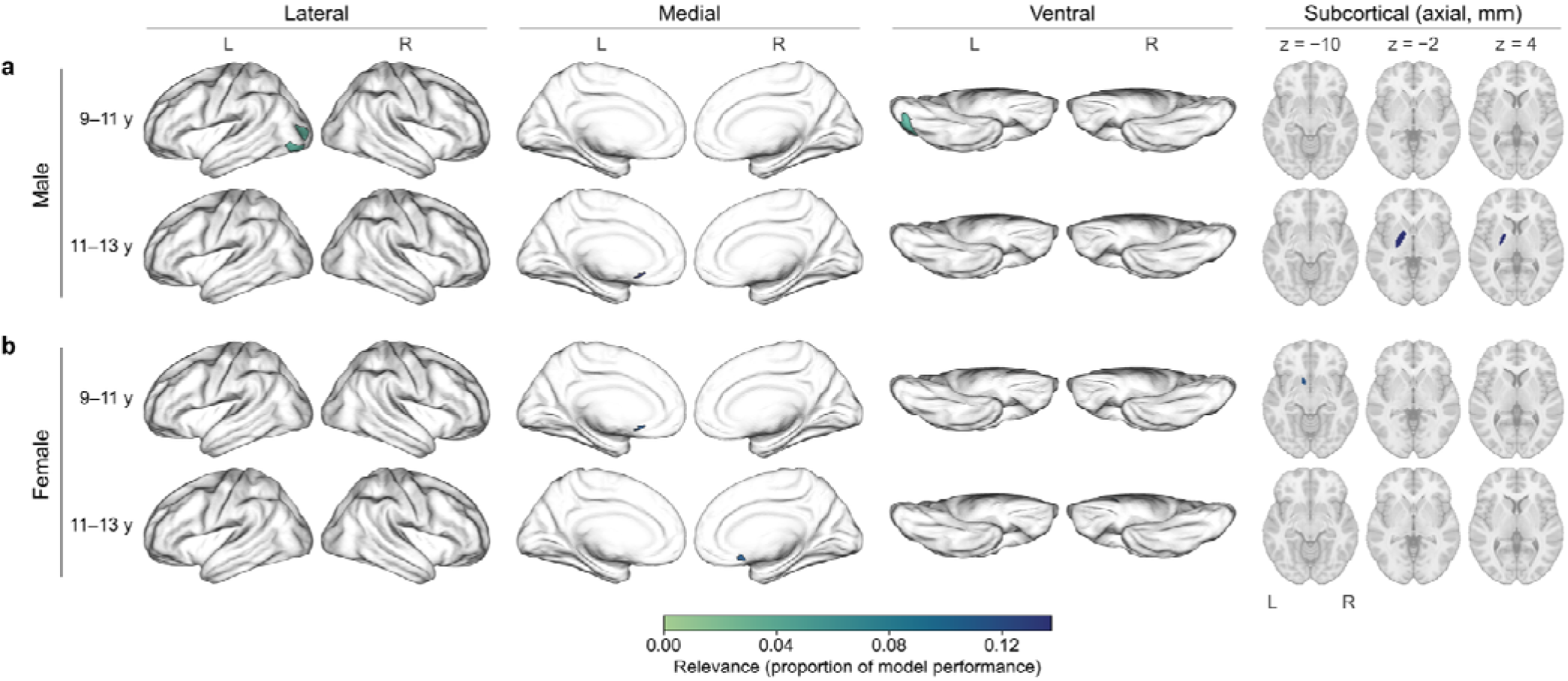
Regions exhibiting individual differences in RNI that were consistently relevant to pubertal testosterone predictions across model iterations. Color-coding indicates the average change in model performance associated with randomizing participant values for each region, across 50 permutations (per region, per cross-validation iteration).

For female youth aged 9-11 years (Figure 5b), individual differences in RNI explained 11.5% of the variance in pubertal DHEA levels (R^2^= 0.115 ± 0.054). The left subcallosal gyrus and left accumbens accounted for 11.5% of model-explained variance in pubertal testosterone. At ages 11-13 years (Figure 5b), individual differences in RNI explained 11.0% of the variance in pubertal DHEA levels (R^2^= 0.110 ± 0.037) driven by the right subcallosal gyrus, which accounted for 11.2% of model-explained variance in pubertal DHEA.

Relevant regions from each model are presented in Figure 5 and Supplementary Table 3.

#### Neurite density (RND)

For male youth aged 9-11 years (Figure 6a), individual differences in RND explained 3.9% of the variance in pubertal testosterone levels (*R*^2^ = 0.039 ± 0.043). Six regions were relevant in all models tested: left inferior part of the precentral sulcus, bilateral middle occipital sulcus and lunatus sulcus, right orbital sulci, right posterior transverse collateral sulcus, and the left inferior occipital gyrus and sulcus. At ages 11-13 years (Figure 6d), the only region relevant in over 85% of models was the forceps major which was relevant in 87% of model iterations.

**Figure 6.**
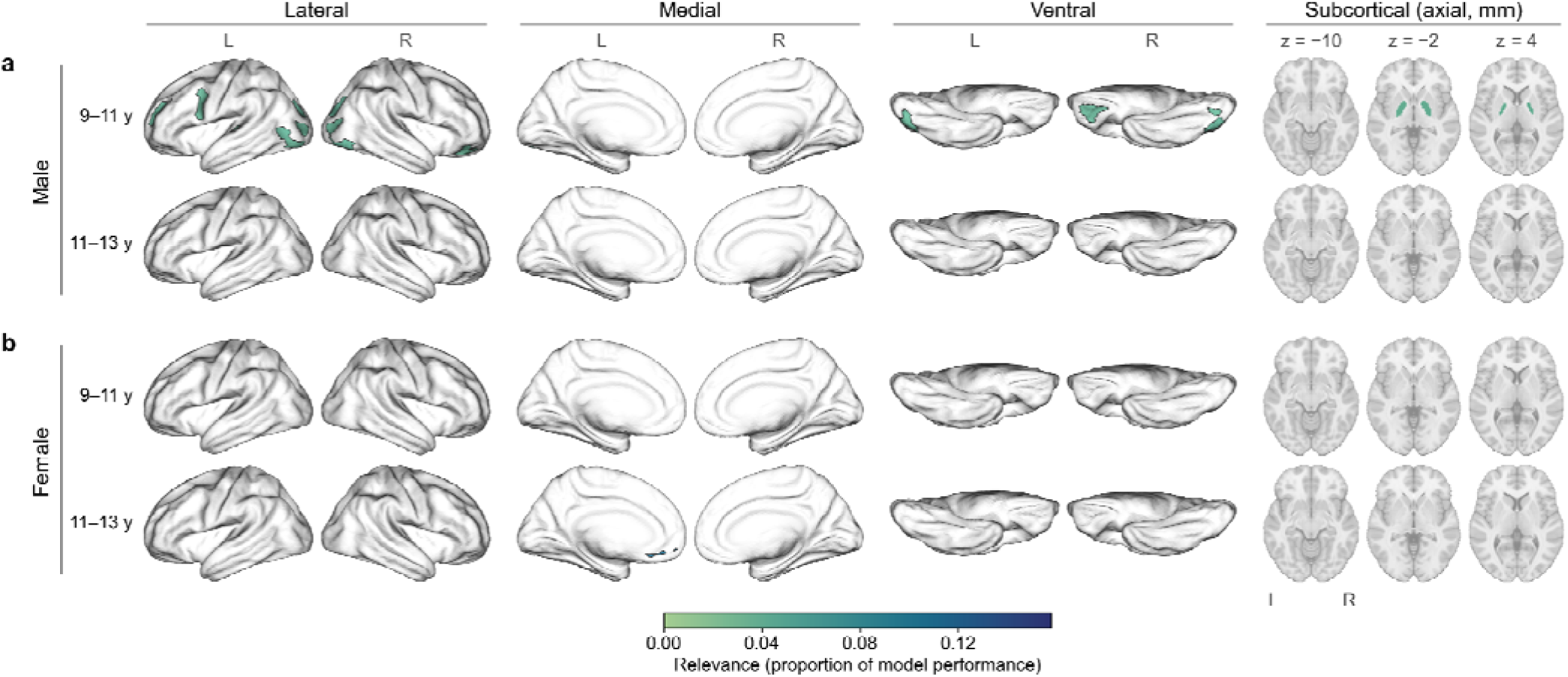
Regions exhibiting individual differences in RND that were consistently relevant to pubertal testosterone predictions across model iterations. Color-coding indicates the average change in model performance associated with randomizing participant values for each region, across 50 permutations (per region, per cross-validation iteration).

For female youth aged 9-11 years (Figure 6a), no regions were relevant in more than 85% of model iterations. At ages 11-13 years, only the left suborbital sulcus was sufficiently relevant, in 92% of models (Figure 6b, arrows). Testosterone accounted for 11% (R^2^) of the variance in RND for this region. No regions were relevant for both males and females at ages 9-11 years or 11-13 years. Relevant regions from all models are presented in Figure 6 and Supplementary Table 4.

#### Microarchitecture related to pubertal estradiol differences

Associations between microarchitecture and estradiol could only be evaluated in the female sample because estradiol was not measured in male youth. The subcallosal gyrus was the only region relevant to pubertal estradiol levels (Figure 7, Supplementary Table 5). At ages 9-11 years (Figure 7a), individual differences in left subcallosal gyrus RNI accounted for 11.1% of model-explained variance in pubertal estradiol levels; at ages 11-13 years, 10.1% of model-explained variance. Individual differences in RND were not relevant to pubertal estradiol predictions at either are range.

**Figure 7.**
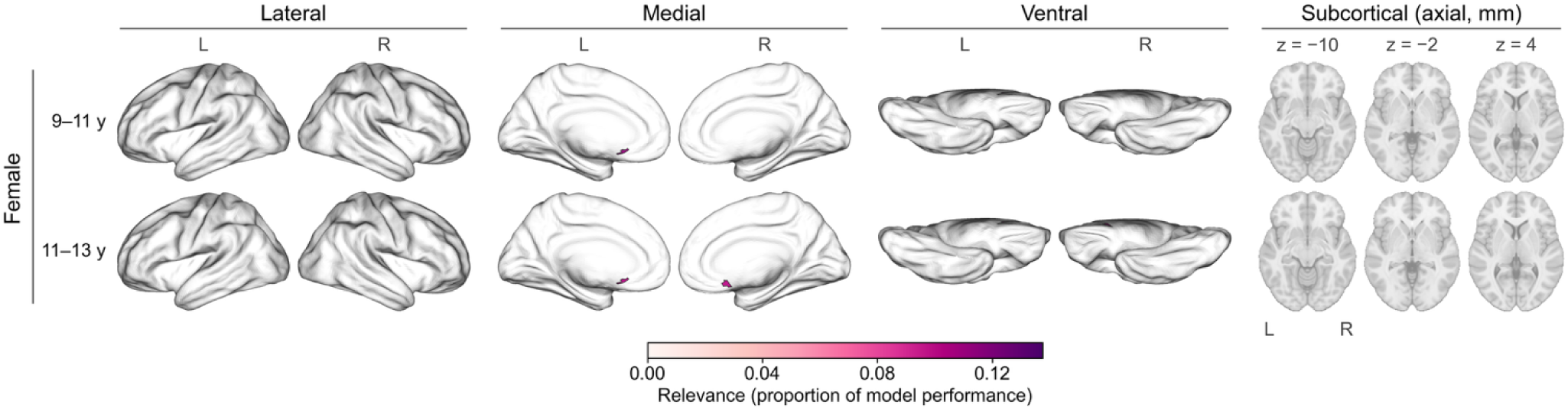
Regions exhibiting individual differences in RNI that were consistently relevant to pubertal estradiol predictions across model iterations. Color-coding indicates the average change in model performance associated with randomizing participant values for each region, across 50 permutations (per region, per cross-validation iteration).

## Discussion

Our sex-stratified predictive modeling framework mapped individual differences in brain microarchitecture to joint information from pubertal hormones and physical development in youth aged 9–13 years of age from the ABCD Study. We found links between pubertal development and regional microarchitecture (neurite and cellular density) that may reflect divergent sex differences in pubertal timing, while elucidating pubertal refinements in reward circuitry. At ages 9-11 years, microarchitecture explained more pubertal hormone variance in female youth (∼11%) than male youth (∼4–6%). By ages 11 to 13 years, however, explained variance in male youth reached comparable levels (∼13–15%) to that of female youth, which remained largely stable (∼11%). At ages 9-11 years, male youth exhibited widespread DHEA-related patterns of both neurite density (RND) and cellular density (RNI). By ages 11–13 years, associations were no longer found between DHEA and neurite density, and relationships between DHEA and cellular density consolidated into fewer, stronger associations. Conversely, at ages 9-11 years, female youth exhibited few DHEA-and testosterone-related microarchitecture associations (both neurite and cellular density), but increased DHEA-related cellular density at ages 11-13 years. Anatomically, hormone-sensitive microarchitecture converged in reward-related regions (e.g., pallidum, subcallosal gyrus, nucleus accumbens), but several regions with known high gonadal hormone receptor expression (e.g., hippocampus, amygdala, cerebellum) did not show significant associations. This exploratory framework revealed complexities in pubertal neuroendocrine development, suggesting differences in trajectories by sex and between microarchitecture properties, providing neurobiological targets for future longitudinal studies.

### Sex differences in hormone-microarchitecture links likely reflect divergent pubertal timing

Microarchitecture explained greater variance in female youth (11-13%) compared to male youth (4-6%) at ages 9-11 years, likely reflecting earlier pubertal timing and related higher salivary hormone levels in female youth (Abreu & Kaiser, 2016). This divergence aligns with the observed two-year lag between sexes in pubertal timing whereby male youth at 11-13 years old reach comparable pubertal stages to female youth at 9-11 years old (Table 3), as microarchitecture-explained variance nearly doubled over this period (i.e., to 9-14%), paralleling increased hormone-PDS correlations. In male youth, the best-performing modes at 11-13 years old were sparser but explained more variance than those at 9-11 years old, which included smaller contributions across more brain areas. This suggests that the pubertal rise in testosterone influences a more specific neuroanatomy. Notably, microarchitecture-explained variance in female youth was stable from ages 9-11 to 11-13 years (11-13%), despite an increased estradiol-PDS correlation, suggesting a persistent sensitivity of brain microarchitecture to hormone shifts into mid-puberty. Further, the plateau in females and decline in males of the number of important testosterone-and DHEA-sensitive microarchitecture regions may reflect the closure of sex-specific sensitive periods during early adolescence, consistent with rodent data showing a window of activity-dependent GABAergic circuit refinement that closes by young adulthood (Caballero et al., 2021). Together, these patterns indicate that pubertal timing shapes hormone-brain interactions differently across sexes and within individuals over time.

### Developmental timing and compartment-specific effects of adrenal and gonadal hormones

In male youth aged 9-11 years, both DHEA and testosterone exhibited more associations with regional neurite density (RND) than cellular density (RNI), suggesting a preferential role in pubertal dendritic and axonal maturation (e.g., myelination, axonal packing). This aligns with animal studies demonstrating testosterone’s influence on dendritic spine density and neurite growth in cortical and limbic development (Chen et al., 2013; Cunningham et al., 2007), while DHEA enhances neurite outgrowth (Lazaridis et al., 2011). However, by ages 11-13 years, the number of regions exhibiting testosterone-and DHEA-related microarchitecture differences decreased dramatically, despite greater explained variance, indicating that organizational effects may be more targeted and pronounced as puberty progresses. In line with this theory, female youth showed fewer regions with testosterone-related associations at either age range for both RNI and RND, compared to their male peers, potentially reflecting fewer, stronger brain-hormone associations because of more advanced puberty. However, we are unable to determine whether the same decrease in number but increase in strength of associations happens in female youth, as the ABCD Study does not include data on youth younger than 9 years old. Across age ranges, female youth showed a similar number of regions exhibiting testosterone-sensitive RNI, but a substantial decrease in regions exhibiting testosterone-sensitive RND (from 4 regions at 9-11 years to only 1 region at 11-13 years). Thus, testosterone-related effects on microarchitecture development in female youth may be more complete by ages 11-13 years.

Conversely, DHEA-sensitive microarchitecture followed a distinct pattern in female youth: more regions exhibited DHEA-sensitive RNI at 11-13 years of age. This may reflect an increasing role for DHEA in cell body development as puberty progresses or DHEA’s function as a precursor for other neuroactive hormones (Zwain & Yen, 1999). To the latter point, DHEA-RNI associations may indirectly reflect downstream estrogenic effects on microarchitecture. While we did measure estradiol in female youth, mean estradiol levels did not differ between the 9-11-and 11-13-year age ranges, and estradiol showed minimal associations with RNI or RND. We suspect that this reflects inherent measurement limitations of quantifying estradiol in this population (e.g., time of sample collection, menstrual cycle phase in postmenarcheal youth). By contrast, DHEA is an upstream adrenal androgen with less acute cyclical variation (Frederiksen et al., 2024; Matchock et al., 2007) that may serve as a more stable proxy for cumulative estrogenic exposure, capturing the sustained developmental influence of estradiol and other neuroactive steroid hormones. The increasing DHEA-RNI associations observed at 11-13 years in female youth may therefore reflect the growing contribution of DHEA-derived estradiol to microarchitecture development as the adrenal and gonadal axes become more active across puberty.

### Pubertal DHEA-related microarchitecture may reflect GABAergic development

Cellular and neurite density (i.e., RNI, RND) in the pallidum were consistently linked to pubertal DHEA and testosterone levels. The pallidum is a central reward processing hub that receives substantial GABAergic inhibitory input from the striatum (Galvan et al., 2005), encodes hedonic reward value, processes risk-reward trade-offs, and regulates decision-making (K. S. Smith et al., 2009a; Soares-Cunha & Heinsbroek, 2023). Given the profound maturation of reward circuitry during adolescence (Galvan, 2010) and GABAergic signaling by both DHEA and testosterone (Wang, 2011), these associations may reflect hormone-mediated refinement of reward processing and motivated behavior during the pubertal transition.

Specifically, DHEA and testosterone may refine this circuitry by moderating its excitatory-inhibitory (E-I) balance. DHEA acts as a GABA_A_ receptor antagonist, increasing neuronal excitability (Paul & Purdy, 1992), while testosterone metabolites act as positive modulators that enhance GABAergic inhibition (Frye et al., 1996). Instead of counteracting each other, DHEA and testosterone may exert opposing effects that drive activity-dependent microstructural development in the pallidum. One hypothesis is that our cortical findings may reflect adolescent E/I balance refinement, as orbitofrontal cellular density was consistently related to individual differences in DHEA levels. Orbitofrontal excitatory-inhibitory balance develops during adolescence as local GABAergic neurons interact with glutamatergic projections (Caballero et al., 2021) and DHEA binds to both (Compagnone & Mellon, 1998). Hence, the observed microarchitecture differences may suggest a mechanism by which adrenarche contributes to the adolescent calibration of cortical excitability.

### Regions with high receptor expression but not linked to pubertal hormones

Contrary to expectations, we did not find hormone-microarchitecture associations in several regions with high gonadal hormone receptor expression, including the hippocampus, amygdala, and cerebellum (Sjöstedt et al., 2020; Zsido et al., 2017). This may be explained by our multivariate approach, which examines multiple relationships across regions simultaneously, and differs from the univariate associations typically reported in the literature. Alternately, these findings may be because we averaged RNI and RND across these regions, which are highly heterogeneous in cytoarchitecture, receptor expression, and developmental profiles (Campbell et al., 2022b; Ojha et al., 2025; Simerly et al., 1990). When microstructural measures are averaged across whole structures, as in our study, localized hormone-microarchitecture relationships are likely obscured, resulting in these null effects. Future investigations should examine more fine-grained differences within hippocampal subfields and amygdala nuclei, especially because both structures project to the nucleus accumbens and ventral pallidum (K. S. Smith et al., 2009b; Yang & Mogenson, 1985) where we did observe robust associations. Pubertal hormones may preferentially influence the integration of these connections within downstream regions rather than the structures themselves.

In the cerebellum, our null findings likely reflect unique challenges for diffusion MRI-based microstructural analyses. The cerebellum features highly specialized layer-specific architecture with perpendicular fiber systems (e.g., radial Purkinje dendrites and tangential parallel fibers in the molecular layer) and exceptionally high densities of parallel fibers (Tamanini et al., 2025). The planar rather than columnar geometry of the Purkinje cell dendrites in the cerebellum, and this region’s perpendicular fiber systems likely violate fundamental RSI model assumptions. Combined with partial volume effects across cerebellar layers with dramatically different cellular properties (sparse molecular layer vs. extremely dense granule cell layer), this diffusion modeling approach may not be well-suited for capturing cerebellar microarchitecture.

### Strengths & Limitations

While the present study leveraged robust covariate regression, sex-stratified analyses, and histologically validated RSI metrics to evaluate hormone-microarchitecture relationships, several limitations remain. First, temporal delays between saliva sample collection and MRI data acquisition ranged from hours to weeks and precluded a precise estimation of hormone states during scans. Further, single measurements cannot account for phasic hormone fluctuations (e.g., diurnal, menstrual), but the additional inclusion of PDS scores in multi-task regression models likely served as a smoothing term for noisier hormone estimates. That is, the greater temporal stability of PDS scores might compensate for greater variability in hormone estimates. Because organizational brain changes during puberty likely reflect tonic increases in hormones, our findings may capture developmentally meaningful associations despite this imprecision. Second, observational data and cross-sectional analyses limits causal inference and hinder our ability to distinguish between lasting organizational changes and transient activational effects. Third, our stringent quality control guidelines for both neuroimaging and endocrine data resulted in substantial attrition and likely introduced selection bias. Combined with the ABCD Study’s sample bias (Compton et al., 2019), our final analytic sample is less racially and socioeconomically diverse than the U.S. population. Future research should collect multiple hormone samples in a narrower window of time that is time-locked with MRI data acquisition and aim to recruit more youth from underrepresented backgrounds, for a more precise and generalizable study of these neuroendocrine mechanisms.

## Conclusion

Our findings demonstrate the importance of sex-stratified, developmentally informed analyses for identifying robust associations between pubertal hormones and brain microarchitecture. By jointly modeling perceived pubertal development and circulating hormone levels, our predictive modeling framework uncovered a few aspects of pubertal neuroendocrine development that warrant future study. First, sex differences in pubertal timing and in the observed microstructural sensitivity to pubertal hormones may, together, suggest that a narrowing developmental window as puberty progresses and may elucidate the emergence of sex differences in adolescent psychopathology. Second, we consistently identified hormone-sensitive microarchitecture in reward processing regions like the pallidum and orbitofrontal cortex that undergo substantial GABAergic refinement during adolescence. Given opposing influences of DHEA and testosterone metabolites on GABAergic neurotransmission, our findings suggest that future work should investigate hormone-mediated calibration excitatory-inhibitory balance within reward circuitry. Microstructural measures in these regions may therefore serve as neurobiological markers of pubertal maturation and could inform our understanding of adolescent-onset mental health conditions, particularly those characterized by disruptions in reward processing and mood regulation. Future research should apply a precision adolescent health framework with repeated hormone measurements, high-resolution neuroimaging, and mental health monitoring to translate this exploratory evidence of hormone-sensitive brain microarchitecture into predictive markers of psychopathology risk.

## Acknowledgments

A special thank you to all of the children and families for their participation in their ABCD Study.

Research described in this article was supported by the National Institutes of Health (KLB: NIMH R00MH135075).

Data used in the preparation of this article were obtained from the Adolescent Brain Cognitive Development^SM^ (ABCD) Study (https://abcdstudy.org), held in the NIMH Data Archive (NDA). This is a multisite, longitudinal study designed to recruit more than 10,000 children aged 9-11 years old and follow them over 10 years into early adulthood. The ABCD Study® is supported by the National Institutes of Health and additional federal partners under award numbers U01DA041048, U01DA050989, U01DA051016, U01DA041022, U01DA051018, U01DA051037, U01DA050987, U01DA041174, U01DA041106, U01DA041117, U01DA041028, U01DA041134, U01DA050988, U01DA051039, U01DA041156, U01DA041025, U01DA041120, U01DA051038, U01DA041148, U01DA041093, U01DA041089, U24DA041123, U24DA041147. A full list of supporters is available at https://abcdstudy.org/federal-partners.html. A listing of participating sites and a complete listing of the study investigators can be found at https://abcdstudy.org/consortium_members/. ABCD consortium investigators designed and implemented the study and/or provided data but did not necessarily participate in the analysis or writing of this report. This manuscript reflects the views of the authors and may not reflect the opinions or views of the NIH or ABCD consortium investigators. The ABCD data repository grows and changes over time. The ABCD data used in this report came from https://dx.doi.org/10.15154/z563-zd24.

## Author Contributions

EC contributed to the writing of the original draft, writing review and editing and contributed to conceptualization of study findings interpretations. ED contributed to writing, review, and editing. MMH contributed to funding acquisition, resources, writing review and editing. KLB conceptualized the work, developed the methodology, conducted the formal analyses, contributed to the visualizations, writing of the original draft, writing review and editing, and project supervision.

