## Supplementary Materials for "Beyond morphometry: Adolescent brain microarchitecture reflects sex differences in pubertal timing and hormones"

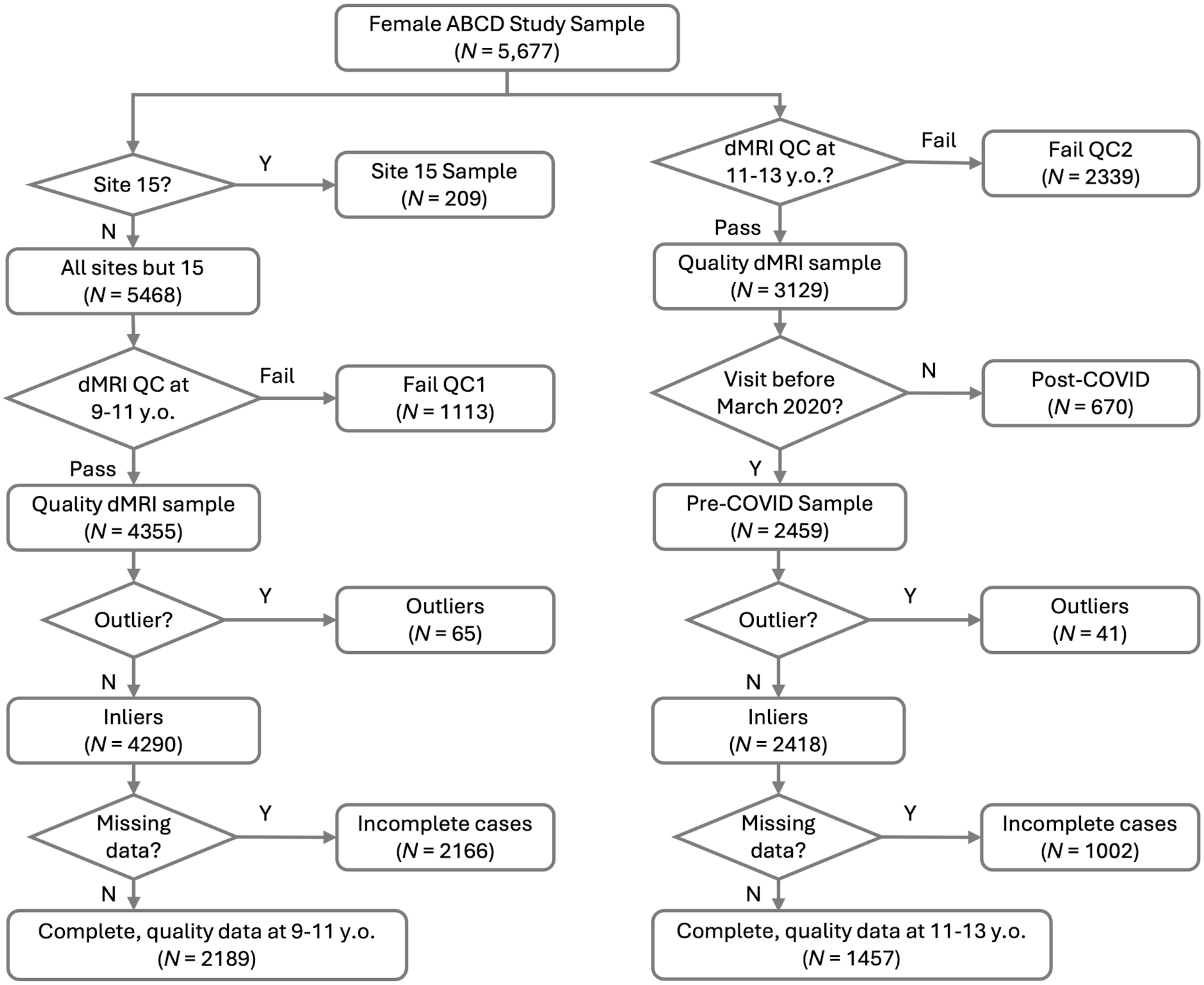


***Supplementary Figure 1. Female participant inclusion flow diagram.***

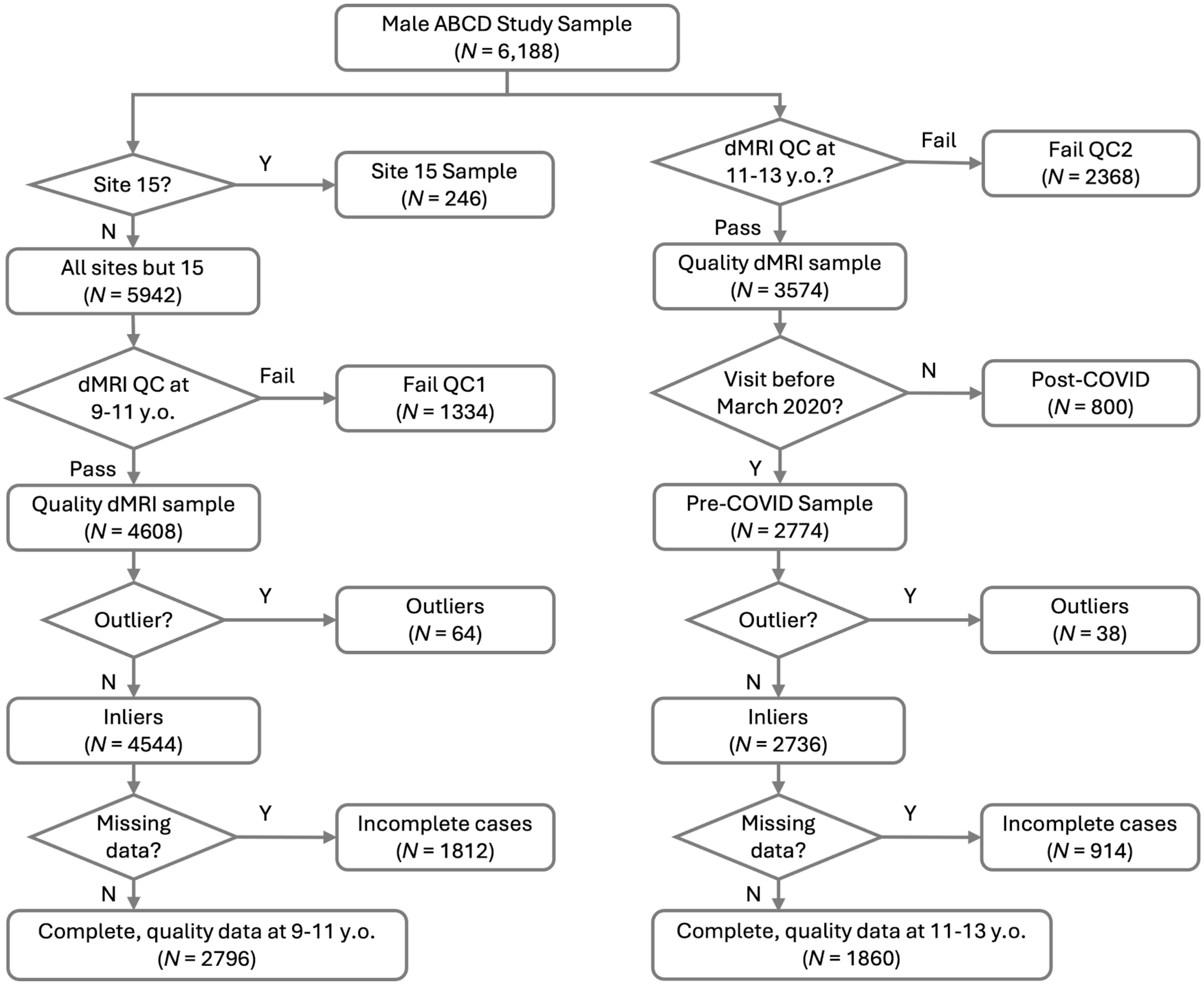


***Supplementary Figure 2. Male participant inclusion flow diagram.***

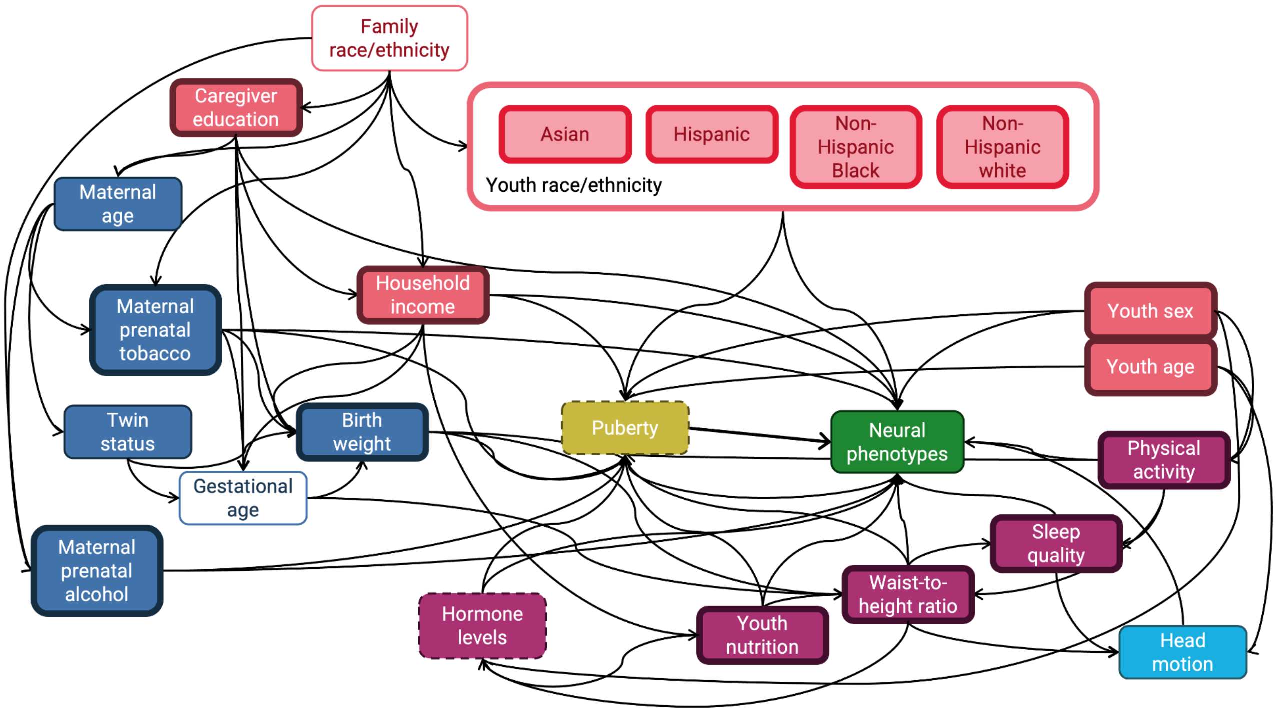


***Supplementary Figure 3. Directed acyclic graph illustrating theory-driven associations between puberty, hormone levels, and neural phenotypes. as well as neuroendocrine confounds.***

Dark blue rectangles denote perinatal variables, pink rectangles denote sociodemographic variables, magenta rectangles indicate physical health variables, light blue square indicates an MRI precision variable that is also on a causal pathway, and unfilled rectangles indicate unmodeled variables in the ABCD Study dataset. Bold outlines indicate the minimally sufficient adjustment set that accounts for the biasing paths identified here (Shrier & Platt, 2008; Textor et al., 2016), while dashed outlines indicate the target variables (i.e., puberty, hormone levels) in the multitask regression model used here. Green rectangle indicates the "outcome"variable, neural phenotypes (i.e., restricted normalized isotropic and directional diffusion, proxies for cellular and neurite density, respectively), while yellow rectangles indicate the "exposure" variables (here: puberty). We acknowledge that many of the key relationships identified here are, in reality, multidirectional (e.g., between brain development, pubertal development, and hormones), but DAG conventions specify unidirectional associations.

***Supplementary Table 1. Feature importance represented by $R^2$ across iterations for models using RNI to predict pubertal DHEA in each sex.***

| | $R^2$ Mean ± Standard Deviation (% relevance) | | | |
|---|---|---|---|---|
| | Males | | Females | |
| Region (hemisphere) | 9-11 years | 11-13 years | 9-11 years | 11-13 years |
| *Subcortical* | | | | |
| Pallidum L | | 0.091 ± 0.056 (100%) | 0.127 ± 0.04 (99%) | |
| Pallidum R | | 0.091 ± 0.057 (97%) | | 0.114 ± 0.034 (99%) |
| Accumbens area L | 0.051 ± 0.035 (88%) | | 0.127 ± 0.04 (100%) | 0.114 ± 0.034 (100%) |
| Accumbens area R | 0.051 ± 0.035 (89%) | | | 0.114 ± 0.034 (100%) |
| *Frontal cortex* | | | | |
| Suborbital sulcus R | 0.048 ± 0.035 (86%) | | | |
| Medial orbital sulcus L | 0.050 ± 0.035 (95%) | | | 0.113 ± 0.034 (95%) |
| Medial orbital sulcus R | 0.050 ± 0.035 (100%) | | | |
| Subcallosal gyrus L | | 0.091 ± 0.056 (100%) | 0.127 ± 0.04 (98%) | 0.114 ± 0.034 (100%) |
| Subcallosal gyrus R | 0.050 ± 0.036 (92%) | | | 0.114 ± 0.034 (100%) |
| *Temporal cortex* | | | | |
| Transverse temporal sulcus L | 0.050 ± 0.035 (95%) | | | |
| *Occipital cortex* | | | | |
| Inferior occipital gyrus and sulcus L | 0.052 ± 0.035 (88%) | | | |
| Inferior occipital gyrus and sulcus R | 0.050 ± 0.035 (94%) | | | |
| Middle occipital sulcus and lunatus sulcus L | 0.050 ± 0.035 (100%) | | | |
| Middle occipital sulcus and lunatus sulcus R | 0.050 ± 0.035 (100%) | | | |
| Anterior occipital sulcus and preoccipital notch L | 0.050 ± 0.035 (96%) | | | |

***Supplementary Table 2. Feature importance represented by $R^2$ across permutations for models using RND to predict DHEA in both sexes.***

| | Mean ± Standard Deviation (% relevance) | | | |
|---|---|---|---|---|
| | Males | | Females | |
| Region (hemisphere) | 9-11 years | 11-13 years | 9-11 years | 11-13 years |
| *Subcortical* | | | | |
| Pallidum R | 0.050 ± 0.036 (100%) | | | |
| *Frontal cortex* | | | | |
| Middle frontal sulcus L | 0.050 ± 0.036 (100%) | | | |
| Middle frontal sulcus R | 0.054 ± 0.034 (87%) | | | |
| Inferior part of the precentral sulcus L | 0.050 ± 0.036 (100%) | | | |
| Inferior part of the precentral sulcus R | 0.052 ± 0.035 (92%) | | | |
| Posterior ramus of the lateral sulcus L | 0.051 ± 0.035 (93%) | | | |
| Orbital sulci R | 0.050 ± 0.036 (100%) | | | |
| Medial orbital sulcus R | 0.052 ± 0.036 (85%) | | | |
| Pericallosal sulcus L | | | 0.128 ± 0.039 (85%) | |
| *Temporal cortex* | | | | |
| Planum temporale L | 0.050 ± 0.036 (100%) | | | |
| Planum temporale R | 0.051 ± 0.035 (91%) | | | |
| Posterior transverse collateral sulcus R | 0.050 ± 0.036 (98%) | | | |
| Transverse temporal sulcus L | 0.050 ± 0.036 (100%) | | 0.127 ± 0.041 (95%) | |
| Transverse temporal sulcus R | 0.051 ± 0.035 (95%) | | | |
| *Occipital cortex* | | | | |
| Anterior occipital sulcus and preoccipital notch L | 0.050 ± 0.036 (98%) | | | |
| Superior occipital sulcus and transverse occipital sulcus L | 0.050 ± 0.036 (99%) | | | |
| Superior occipital sulcus and transverse occipital sulcus R | 0.049 ± 0.035 (87%) | | | |
| Middle occipital sulcus and lunatus sulcus L | 0.050 ± 0.036 (100%) | | | |

***Supplementary Table 2. Feature importance represented by $R^2$ across permutations for models using RND to predict DHEA in both sexes.***

| | | |
|---|---|---|
| Middle occipital sulcus and lunatus sulcus R | 0.050 ± 0.036 (96%) | 0.114 ± 0.037 (91%) |
| Inferior occipital gyrus and sulcus L | 0.050 ± 0.036 (100%) | |
| Inferior occipital gyrus and sulcus R | 0.050 ± 0.036 (100%) | |
| *White matter* | | |
| Cingulate cingulum L | | 0.109 ± 0.036 (88%) |

***Supplementary Table 3. Feature importance represented by $R^2$ across permutations for models using RNI to predict testosterone in both sexes. Only regions that were relevant in at over 85% of model iterations are reported.***

| | Mean ± Standard Deviation (% relevance) | | | |
|---|---|---|---|---|
| | Males | | Females | |
| Region (hemisphere) | 9-11 years | 11-13 years | 9-11 years | 11-13 years |
| *Subcortical* | | | | |
| Accumbens area L | | | 0.115 ± 0.053 (98%) | |
| Pallidum L | | 0.137 ± 0.042 (97%) | | |
| *Frontal cortex* | | | | |
| Subcallosal gyrus L | | 0.136 ± 0.043 (99%) | 0.115 ± 0.053 (100%) | |
| Subcallosal gyrus R | | | | 0.112 ± 0.037 (95%) |
| *Occipital cortex* | | | | |
| Middle occipital sulcus and lunatus sulcus L | 0.040 ± 0.043 (99%) | | | |
| Inferior occipital gyrus and sulcus L | 0.043 ± 0.042 (91%) | | | |

***Supplementary Table 4. Feature importance represented by $R^2$ across permutations for models using RND to predict testosterone in both sexes. Only regions that were relevant in at over 85% of model iterations are reported.***

| Region (hemisphere) | Mean ± Standard Deviation (% relevance) | | | |
|---|---|---|---|---|
| | Males | | Females | |
| | 9-11 years | 11-13 years | 9-11 years | 11-13 years |
| *Subcortical* | | | | |
| Pallidum L | 0.038 ± 0.044 (94%) | | | |
| Pallidum R | 0.039 ± 0.044 (97%) | | | |
| *Frontal cortex* | | | | |
| Middle frontal sulcus L | 0.040 ± 0.042 (92%) | | | |
| Inferior part of the precentral sulcus L | 0.039 ± 0.043 (100%) | | | |
| Orbital sulci R | 0.039 ± 0.043 (100%) | | | |
| Suborbital sulcus L | | | | 0.11 ± 0.036 (92%) |
| *Temporal cortex* | | | | |
| Transverse temporal sulcus L | 0.039 ± 0.043 (98%) | | | |
| Posterior transverse collateral Sulcus R | 0.039 ± 0.043 (100%) | | | |
| *Occipital cortex* | | | | |
| Anterior occipital sulcus and preoccipital notch L | 0.041 ± 0.041 (94%) | | | |
| Superior occipital sulcus and Transverse occipital sulcus L | 0.041 ± 0.042 (95%) | | | |
| Superior occipital sulcus and Transverse occipital sulcus R | 0.041 ± 0.041 (86%) | | | |
| Middle occipital sulcus and lunatus sulcus L | 0.039 ± 0.043 (100%) | | | |
| Middle occipital sulcus and lunatus sulcus R | 0.039 ± 0.043 (100%) | | | |
| Inferior occipital gyrus and sulcus R | 0.042 ± 0.041 (92%) | | | |
| Inferior occipital gyrus and sulcus L | 0.039 ± 0.043 (100%) | | | |
| *White matter* | | | | |
| Forceps major | | 0.158 ± 0.028 (87%) | | |

***Supplementary Table 5. Feature importance represented by $R^2$ across permutations for models using RNI to predict estradiol in female youth.***

| Region (hemisphere) | Mean ± Standard Deviation (% relevance) | |
|---|---|---|
| | Ages 9-11 years | Ages 11-13 years |
| Subcallosal gyrus L | 0.111 ± 0.045 (100%) | 0.101 ± 0.037 (100%) |
| Subcallosal gyrus R | | 0.101 ± 0.037 (100%) |
